# High-multiplexity imaging reveals that temporal-to-spatial patterning of embryonic structures can involve active transformation of temporal information rather than direct mapping

**DOI:** 10.64898/2026.05.12.724534

**Authors:** Jimena Garcia-Guillen, Mahla Ahmadi, Theophilus Frimpong, Iris Seaman, Kimberly Pacheco, Iris Ambuehl, Christine Mau, Gifty Duah, Tamer Oraby, Ezzat El-Sherif

**Affiliations:** School of Integrative Biological and Chemical Sciences (SIBCS), University of Texas Rio Grande Valley (UTRGV), Edinburg, TX 78539, USA; School of Mathematical and Statistical Sciences, University of Texas Rio Grande Valley (UTRGV), Edinburg, TX 78539, USA; Department of Computer Science, University of Texas Rio Grande Valley (UTRGV), Edinburg, TX 78539, USA; Department of Evolutionary Developmental Genetics, Georg-August-University Göttingen, 37077 Göttingen, Germany

## Abstract

How spatial patterns arise during embryonic development is classically explained by the French Flag model, in which cells acquire positional identities by interpreting morphogen concentration thresholds. However, in many developmental systems, spatial patterns instead emerge progressively through temporal programs of gene expression that are transformed into spatial organization. For example, in the short-germ insect *Tribolium castaneum*, both periodic pair-rule gene expressions that generate body segments and non-periodic gap gene expressions that establish regional identities arise sequentially at the posterior and propagate anteriorly in waves across the developing embryo. Understanding how such temporal gene expression programs are translated into spatial patterns remains a major challenge. To address this problem, we developed a sequential multiplexed imaging strategy based on hybridization chain reaction (HCR), enabling visualization of up to ten anterior-posterior (AP) patterning genes within the same *Tribolium* embryo. By combining this approach with intronic-exonic labeling, we established a framework to infer gene expression dynamics and propagatory behavior during AP patterning. Using this framework, we show that gap gene expression domains remain dynamic and continue to propagate during tissue elongation, indicating that spatial patterns are actively remodeled throughout development. We then directly compared temporal gene activation at the posterior with the resulting spatial organization of pair-rule and gap genes. Surprisingly, while primary pair-rule genes preserve their temporal phase relationships in space, gap genes do not. Instead, the relative positioning of gap gene domains progressively changes as they move anteriorly, indicating that the final spatial organization of gap genes is actively reshaped during propagation rather than being directly inherited from the initial temporal sequence. The continued propagatory behavior of gap gene domains suggests that such reshaping could arise through differential propagation dynamics between genes and/or through progressive reconfiguration of underlying gene regulatory interactions during pattern formation. Together, these findings reveal that temporal-to-spatial patterning can involve active transformation of temporal information rather than a simple mapping from time into space.

## Introduction

Spatial patterning during embryonic development depends on the coordinated regulation of gene expression across body axes and tissues. Classically, this process has been explained by the French Flag model [1], [2], in which cells acquire positional identities by interpreting morphogen concentration thresholds (Fig. 1A). In this framework, spatial domains arise through local responses to graded signaling inputs, allowing patterns to be specified simultaneously across a tissue. This view has been strongly shaped by studies of anterior-posterior (AP) [3]–[6] and dorsal-ventral (DV) [4], [7]–[9] patterning in *Drosophila melanogaster*.

**Figure 1.**
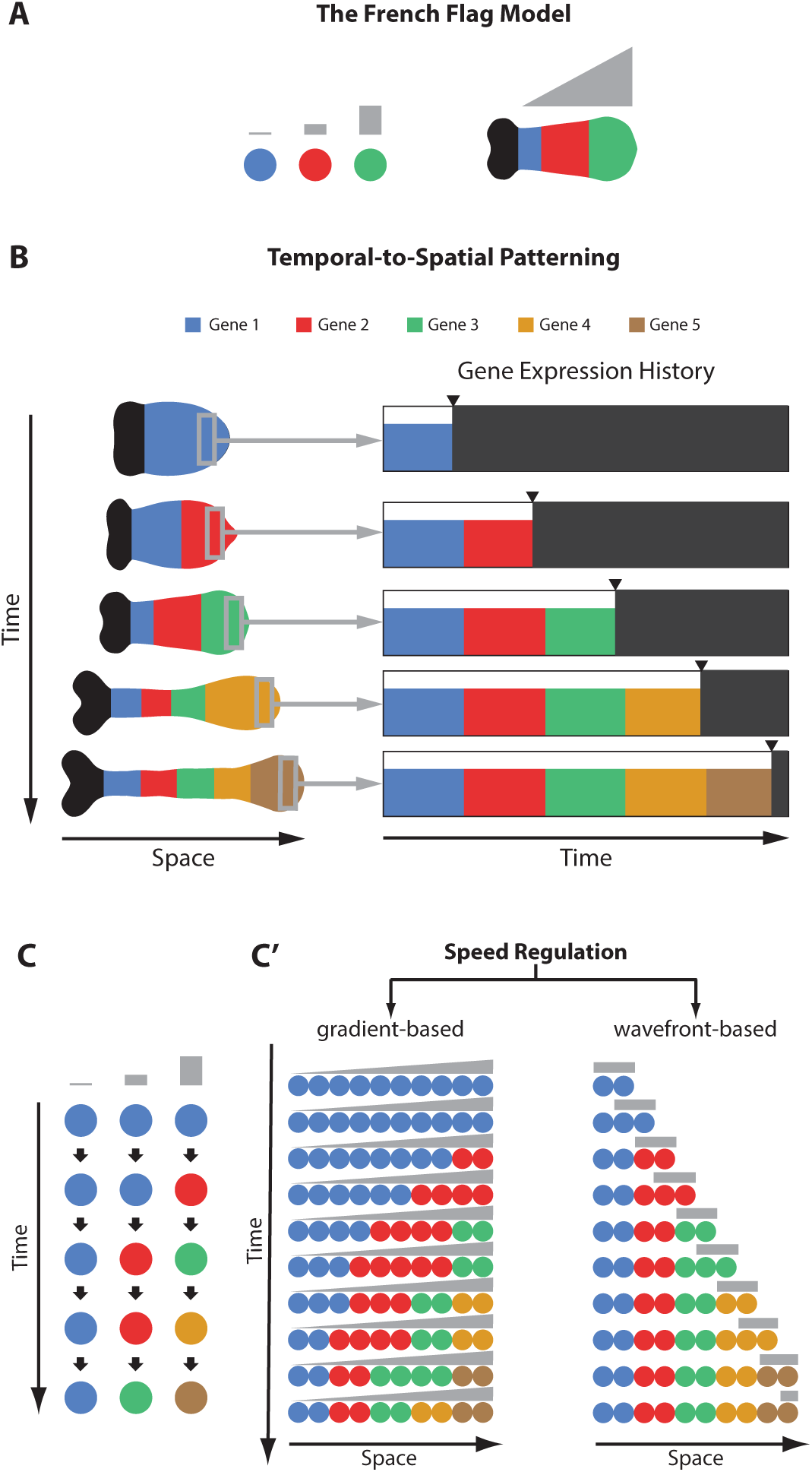
Conceptual frameworks for embryonic spatial patterning. (**A) The French Flag model.** Spatial patterns arise through local interpretation of concentration thresholds of a morphogen gradient (shown in grey), allowing positional identities (shown in different colors, each representing the expression of a different gene) to be specified simultaneously across the tissue. In the illustrated example (left), low concentrations of the morphogen gradient activate a hypothetical blue gene, intermediate concentrations activate a red gene, and high concentrations activate a green gene. The resulting spatial organization along a hypothetical insect AP axis is shown on the right, patterned by a posterior-to-anterior morphogen gradient (posterior to the left). (**B) Temporal-to-spatial patterning:** Spatial organization emerges through the progressive unfolding of a sequential temporal gene expression program. The scheme is illustrated using a hypothetical example of insect AP patterning (posterior to the right). In this example, genes specifying AP identities are sequentially activated through a temporal program operating at the posterior end of the embryo (left; activation sequence from gene 1 to gene 5). As development proceeds, this temporal sequence is progressively translated into a spatial pattern along the AP axis. (**C) The Speed Regulation model:** A possible mechanism for temporal-to-spatial patterning. Sequential gene activation is controlled by a morphogen gradient (shown in grey) acting as a “speed regulator,” which modulates the rate at which cells progress through a temporal sequence of gene expression states (represented in different colors). Cells exposed to high levels of the speed regulator transition rapidly through the sequence, whereas cells exposed to lower levels progress more slowly. **(C’) Modes of Speed Regulation.** <u>In the gradient-based mode (left)</u>, the speed regulator forms a stationary gradient across the tissue. Because cells at different positions progress through the temporal sequence at different rates, propagating waves of gene expression emerge and move from regions of high speed regulator concentration toward regions of lower concentration. <u>In the</u> <u>wavefront-based mode (right)</u>, the speed regulator instead retracts across the tissue over time, and so effectively the speed gradient is acting as a retracting ‘wavefront’. Cells posterior to the retracting wavefront continue progressing dynamically through the temporal sequence, whereas cells anterior to the wavefront become stabilized in their current regulatory state. In this way, the moving wavefront progressively converts temporal progression into stable spatial domains.

However, not all developmental systems conform to this static, threshold-based view. It has long been recognized that certain processes, such as neurogenesis, involve temporal sequences of gene expression that guide cell fate decisions [10]–[15]. In these systems, however, temporal progression is not directly reflected in spatial organization, as cells are redistributed by morphogenetic movements [15]. In contrast, recent work has identified a mode of pattern formation, termed “temporal-to-spatial patterning” (Fig. 1B), in which sequential gene activation events are converted into spatially organized patterns [5], [6], [15]–[24]. Prominent examples include vertebrate segmentation (somitogenesis) [25]–[27], where oscillatory gene expression generates periodic spatial segments, and the sequential activation of Hox genes [28], [29], which defines axial identities along the body axis. Similarly, patterning of the vertebrate neural tube involves the integration of temporal gene activation with morphogen gradients to establish spatial domains along the dorsal-ventral axis [17], [18]. In insects, temporal-to-spatial patterning is prominently observed during AP patterning in *Tribolium castaneum*, where sequential activation of gap and pair-rule genes gives rise to ordered axial domains and segmental patterns [16], [19], [20], [30]. Related propagatory and sequential patterning dynamics have also been described in other insects and arthropods [31]–[37], suggesting that temporal-to-spatial patterning represents a broadly conserved strategy for organizing embryonic tissues.

To account for this mode of pattern formation, we previously proposed the Speed Regulation model [16], [19], [20] (Fig. 1C,C’). In this framework, a morphogen gradient functions as a “speed regulator,” controlling the rate at which cells progress through sequential gene expression states (Fig. 1C). In the gradient-based mode of the model (Fig. 1C’, left panel), cells exposed to high levels of the speed regulator progress rapidly through the temporal sequence, whereas cells exposed to lower levels progress more slowly. As a result, neighboring cells occupy different states of the temporal program depending on their position along the gradient. This generates propagating waves of gene expression that emerge near the signaling center, where the speed regulator concentration is highest, and progressively move toward regions of lower concentration. Importantly, in this mode, spatial patterns arise continuously from differences in progression rates across the tissue rather than from direct threshold specification. This gradient-based mode is particularly suited for patterning non-elongating tissues, where the morphogen gradient is typically static or non-retracting. In the wavefront-based mode of the model (Fig. 1C’, right panel), the speed regulator instead retracts across the tissue as development proceeds. Cells located posterior to the retracting wavefront continue to progress dynamically through the temporal sequence, whereas cells anterior to the wavefront become stabilized in their current state. In this way, the moving wavefront progressively converts temporal progression into stable spatial domains. This wavefront-based mode is particularly suited for elongating tissues, where the morphogen gradient itself retracts as the tissue grows. Thus, in both modes, spatial patterns emerge from temporal gene expression programs, but differ in how these dynamics are translated into stable spatial organization.

The short-germ insect *Tribolium castaneum* provides a particularly suitable system for studying this process. In contrast to long-germ insects such as *Drosophila melanogaster* [38], where most gap and pair-rule gene expression domains arise more or less simultaneously across the blastoderm, AP patterning in *Tribolium* proceeds progressively from anterior to posterior. In *Tribolium*, both pair-rule genes, which mediate segmentation, and gap genes, which mediate AP regionalization, are sequentially activated in the posterior region of the embryo and subsequently translated into spatial patterns along the AP axis [6]. Previous work showed that a posterior Wnt/Caudal gradient modulates both pair-rule oscillations and gap gene dynamics, producing predictable changes in spatiotemporal patterning consistent with the Speed Regulation framework [16], [19], [20].

Importantly, although *Drosophila* patterning is often described in terms of simultaneous spatial specification, both gap and pair-rule genes nevertheless exhibit limited propagatory dynamics [22], [39]–[41], potentially reflecting an evolutionary vestige of an ancestral temporal-to-spatial patterning mechanism. Similarly, DV patterning in *Drosophila*, long considered a paradigmatic example of threshold-based morphogen interpretation, has increasingly been shown to involve sequential and dynamic regulatory processes, suggesting that elements of temporal-to-spatial patterning may operate there as well [42], [43].

A central implication of temporal-to-spatial patterning is that pattern formation depends on the coordinated dynamics of entire gene regulatory networks (GRNs), rather than solely on local threshold responses of individual genes. In models such as the Speed Regulation model, morphogens are not proposed to directly specify spatial domains through concentration thresholds. Instead, the primary role of the morphogen is to modulate the timing of an underlying temporal GRN that mediates sequential gene activation. Spatial patterns therefore emerge from the progression of network states over time and their transformation into spatial organization across the tissue. This shifts the focus of the problem from isolated morphogen-gene interactions toward understanding how the dynamics of entire GRNs are regulated and progressively reshaped during development. Consequently, understanding temporal-to-spatial patterning requires simultaneous analysis of multiple genes and their dynamic relationships within the same developmental context, rather than examining individual genes independently.

However, conventional fluorescence imaging typically permits visualization of only a small number of genes simultaneously due to spectral overlap [44]. As a result, GRN-level analyses often require integration across multiple embryos, obscuring direct spatial and temporal relationships between genes within the same embryo.

Furthermore, temporal-to-spatial patterning, including the Speed Regulation model, fundamentally depends on spatiotemporal gene expression dynamics. In systems exhibiting temporal-to-spatial patterning, sequential gene activation is typically accompanied by propagatory behavior of gene expression domains, in which waves of gene expression emerge near a signaling center and progressively move across the tissue as temporal information is transformed into spatial organization [27], [45]–[47]. Understanding such dynamics therefore requires approaches capable of simultaneously visualizing multiple genes together with measurements that capture their propagatory behavior across developing tissues. While live imaging provides a natural approach to studying developmental dynamics, current approaches remain limited in multiplexity [48], [49]. An alternative strategy exploits the temporal relationship between intronic and exonic transcriptional signals [50], [51]. Intronic signal reflects nascent transcription, whereas exonic signal reflects accumulated mRNA. During propagation of gene expression domains, the temporal offset between these signals generates characteristic spatial phase shifts between intronic and exonic expression domains, providing a proxy for gene expression dynamics in fixed embryos. Importantly, extending this strategy to GRN-level analysis requires simultaneous visualization of both intronic and exonic signals for multiple genes, substantially increasing multiplexing demands.

To address these challenges, we developed a sequential multiplexed imaging strategy based on hybridization chain reaction (HCR) [52], enabling repeated rounds of imaging within the same *Tribolium* embryo (Fig. 2). This approach is conceptually aligned with emerging high-multiplex imaging and spatial genomics technologies that seek to resolve complex molecular organization directly within intact tissues and embryos [53]–[57]. By combining sequential imaging with intronic-exonic labeling, this approach enables simultaneous analysis of multiple genes, their spatial relationships, and their propagatory dynamics within a shared developmental context.

**Figure 2.**
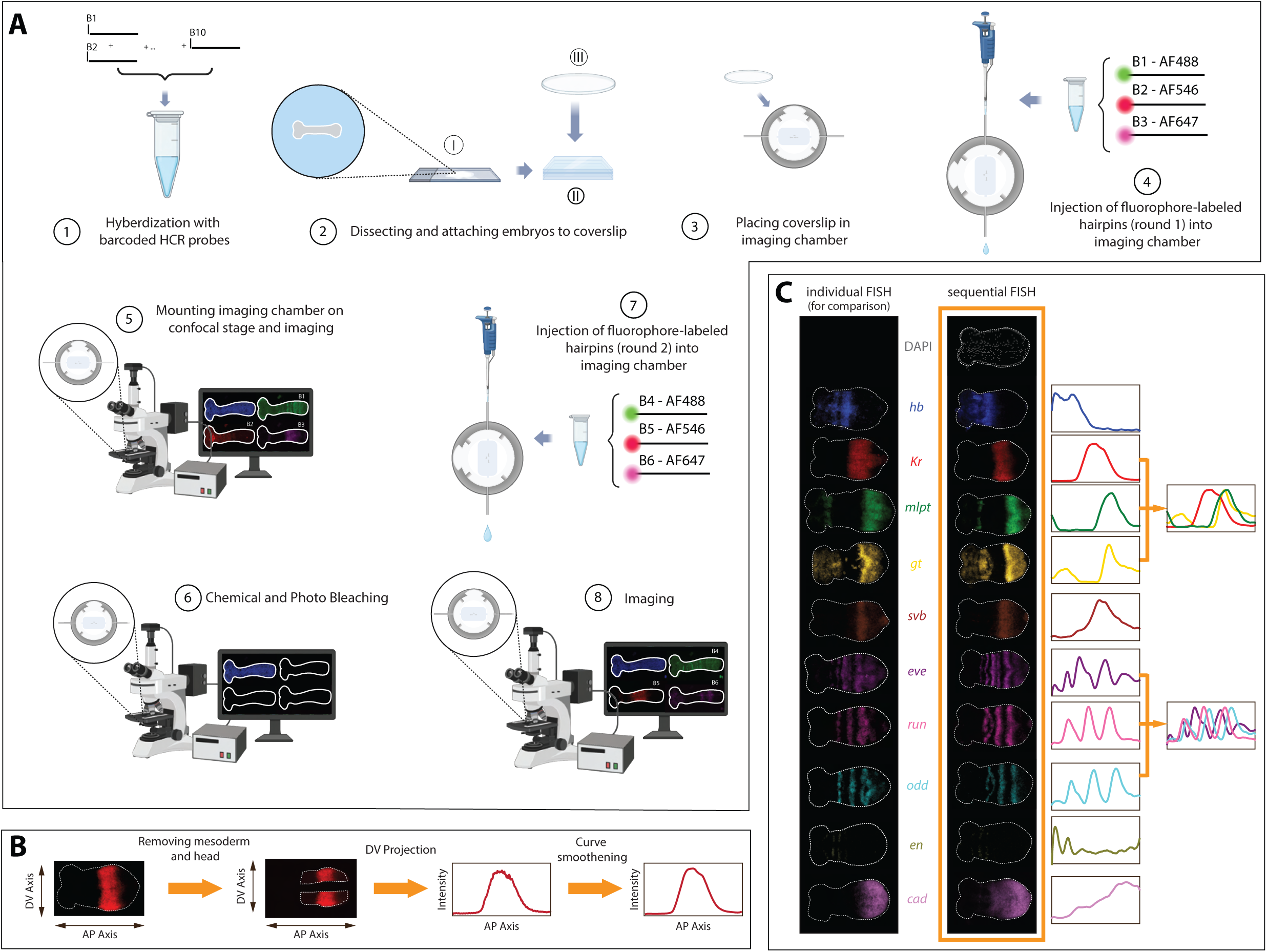
Sequential HCR imaging for high-multiplex gene expression analysis. **(A) Sequential imaging workflow. Step 1**: First, embryos were incubated with HCR probe sets targeting specific transcripts. Each probe set was uniquely barcoded (B1 to B10). **Step 2:** Embryos were then dissected on a microscope slide (I) under a dissecting microscope. The dissected embryos were subsequently transferred onto a PBS agarose gel pad (II) and then gently mounted onto chrome alum gelatin adhesive-coated circular coverslips (III). **Step 3:** The coverslip containing embryos was placed into Bioptechs FCS2 imaging chamber. **Step 4:** Prior to imaging, embryos were incubated in amplification buffer containing fluorophore-labeled HCR hairpin amplifiers, complementary to used barcodes. In the first imaging round, amplifier systems B1, B2, and B3 were paired with Alexa Fluor 488, 546, and 647 fluorophores, respectively, enabling multiplexed visualization of three distinct target transcripts within the same embryo. **Step 5:** Following HCR amplification, embryos were imaged using confocal microscopy to visualize the first set of spatial gene expression patterns (corresponding to barcodes B1 to B3). **Step 6:** Fluorescent signal was subsequently removed through combined chemical bleaching and photobleaching to eliminate residual fluorescence and prevent signal carryover between imaging rounds. **Step 7:** After bleaching, embryos were re-incubated with amplification buffer containing a new set of fluorophore-labeled hairpin amplifiers (e.g., B4-B6) and re-imaged. **Step 8 and beyond:** This sequential cycle of amplification, imaging, bleaching, and re-staining was repeated multiple times, allowing visualization of up to ten genes within the same embryo. **(B) Image processing and quantitative analysis workflow.** Following imaging, mesodermal regions were computationally removed. Fluorescent signal intensity along the DV axis was projected onto the AP axis to generate one-dimensional spatial expression profiles. Curves were then smoothed to facilitate quantitative comparison of gene expression patterns across embryos and developmental stages. **(C) Validation of high multiplexity sequential FISH.** Comparison between sequential imaging and conventional individual HCR FISH stainings confirmed that the sequential imaging protocol faithfully reproduces expected spatial gene expression patterns. Successful validation was obtained for the AP patterning genes *hb, Kr, mlpt, gt, svb, eve, run, odd, en,* and *cad*, demonstrating reliable multiplex visualization of these genes within the same embryo.

Using this framework, we systematically reconstructed both the temporal sequence of AP patterning genes at the posterior signaling center and their resulting spatial organization along the embryo axis. By combining sequential multiplex imaging with intronic-exonic labeling, we were able to simultaneously visualize multiple gap and pair-rule genes, quantify their spatial relationships within the same embryo, and infer aspects of their dynamic propagation during development. This revealed that gap gene expression domains continue to propagate during tissue elongation, demonstrating that AP patterning remains dynamically remodeled even during later developmental stages rather than becoming spatially fixed once domains are established.

We then directly compared the temporal sequence of gene activation at the posterior with the resulting spatial organization of gene expression domains along the embryo axis. This comparison revealed a striking distinction between the two major AP patterning systems analyzed here. Primary pair-rule genes largely preserve their temporal phase relationships in space, consistent with a relatively faithful temporal-to-spatial transformation. In contrast, gap genes exhibit progressive, gene-specific changes in relative spatial positioning as their domains propagate anteriorly. Thus, unlike pair-rule genes, the final spatial organization of gap genes is not simply inherited from the initial temporal sequence generated at the posterior.

Importantly, the observation that gap gene domains continue to propagate provides a potential explanation for this discrepancy. One possibility is that different gap genes propagate with different effective dynamics or velocities, progressively reshaping their relative spatial relationships during tissue elongation. Alternatively, these observations may reflect continuous reconfiguration of the underlying GRNs during propagation, such that the regulatory logic governing spatial organization changes across the tissue. More broadly, our results suggest that temporal-to-spatial patterning cannot always be understood as a simple spatial freezing of an initial temporal program, but may instead involve continuous remodeling of gene expression relationships during pattern propagation.

## Results

### 1. Sequential imaging enables multiplexed analysis of gene expressions within single embryos

To investigate gene expression dynamics across multiple regulatory genes, we implemented a sequential imaging protocol based on HCR (Fig. 2). This approach was designed to allow high-resolution visualization of multiple gene expression patterns within a single biological sample.

Probe sets targeting ten genes involved in AP patterning in the early *Tribolium* embryo (specifically, *hunchback* (*hb*), *Krüppel* (*Kr*), *mille-pattes* (*mlpt*), *giant* (*gt*), *shavenbaby* (*svb*), *even-skipped* (*eve*), *engrailed* (*en*), *caudal* (*cad*), *odd-skipped* (*odd*), and *runt* (*run*)) were first hybridized simultaneously, with each gene assigned a unique HCR barcode, B1 to B10, prior to imaging (Fig. 2A, Step 1). Embryos were then immobilized on slides (Fig. 2A, Step 2) and mounted within an imaging chamber (Fig. 2A, Step 3), allowing stable positioning and repeated reagent exchange during sequential imaging. Sequential imaging was performed through iterative rounds of fluorescent amplification, imaging, and signal bleaching. In each round, two to three barcodes were visualized using complementary HCR hairpin amplifiers carrying distinct fluorophores, for example B1 with AF488, B2 with AF546, and B3 with AF647 (Fig. 2A, Step 4), followed by confocal imaging (Fig. 2A, Step 5). After imaging, fluorescence signals were removed by chemical and photobleaching (Fig. 2A, Step 6). A new set of fluorescent hairpin amplifiers complementary to additional barcodes was then introduced, and the cycle was repeated until all target genes were visualized (Fig. 2A, Steps 7 and 8; see Methods for details).

For quantitative analysis, gene expression was measured along the AP axis within the ectoderm. Because mesodermal expression of some AP genes is delayed relative to ectodermal expression, mesodermal regions were excluded to ensure consistent spatial comparison (Fig. 2B). One-dimensional expression profiles were generated for each gene by projecting signals along the dorsal-ventral (DV) axis and were subsequently smoothened. These profiles yield quantitative representations of the spatial distribution, relative positioning, and overlap of gene expression domains across developmental stages (Fig. 2B).

Using this approach, we visualized the expression of ten AP patterning genes within individual embryos (Fig. 2C; embryos panel to the right). For validation, expression patterns obtained by sequential imaging were compared with conventional single-gene FISH, confirming that the sequential protocol faithfully reproduces expected patterns (Fig. 2C; embryos panel to the left). In addition, we generated corresponding expression profiles for multiple genes, allowing direct comparison of their relative positions (gene expression plots in Fig. 2C).

Together, these results establish a sequential imaging framework capable of producing multiplexed, spatially resolved dataset of gene expressions within single embryos. This framework enables direct comparison between genes within the same spatial context and provides the experimental foundation for analyzing temporal-to-spatial patterning and dynamic gene expression processes.

### 2. Temporal staging framework for AP patterning dynamics

A central goal of this study is to compare temporal gene activation at the posterior signaling center with the resulting spatial organization of gene expression domains along the AP axis. Achieving this requires accurate temporal alignment of embryos across developmental progression. To establish such a framework, we used the dynamic expression patterns of *eve* and *en* as temporal landmarks during AP patterning (Fig. 3A). In *Tribolium*, *eve* is expressed in oscillatory waves at the posterior signaling center, with each oscillatory cycle corresponding to the formation of a pair-rule stripe that ultimately contributes to two segmental units along the AP axis. Each oscillatory cycle progresses through a reproducible sequence of expression states. We therefore subdivided each cycle into three successive stages: initial appearance of *eve* expression at the posterior (stage X.1), expansion of the expression domain (stage X.2), and posterior clearance of expression (stage X.3), where X indicates the oscillation cycle number. These stereotyped dynamics provide a reproducible readout of temporal phase during AP patterning.

**Figure 3.**
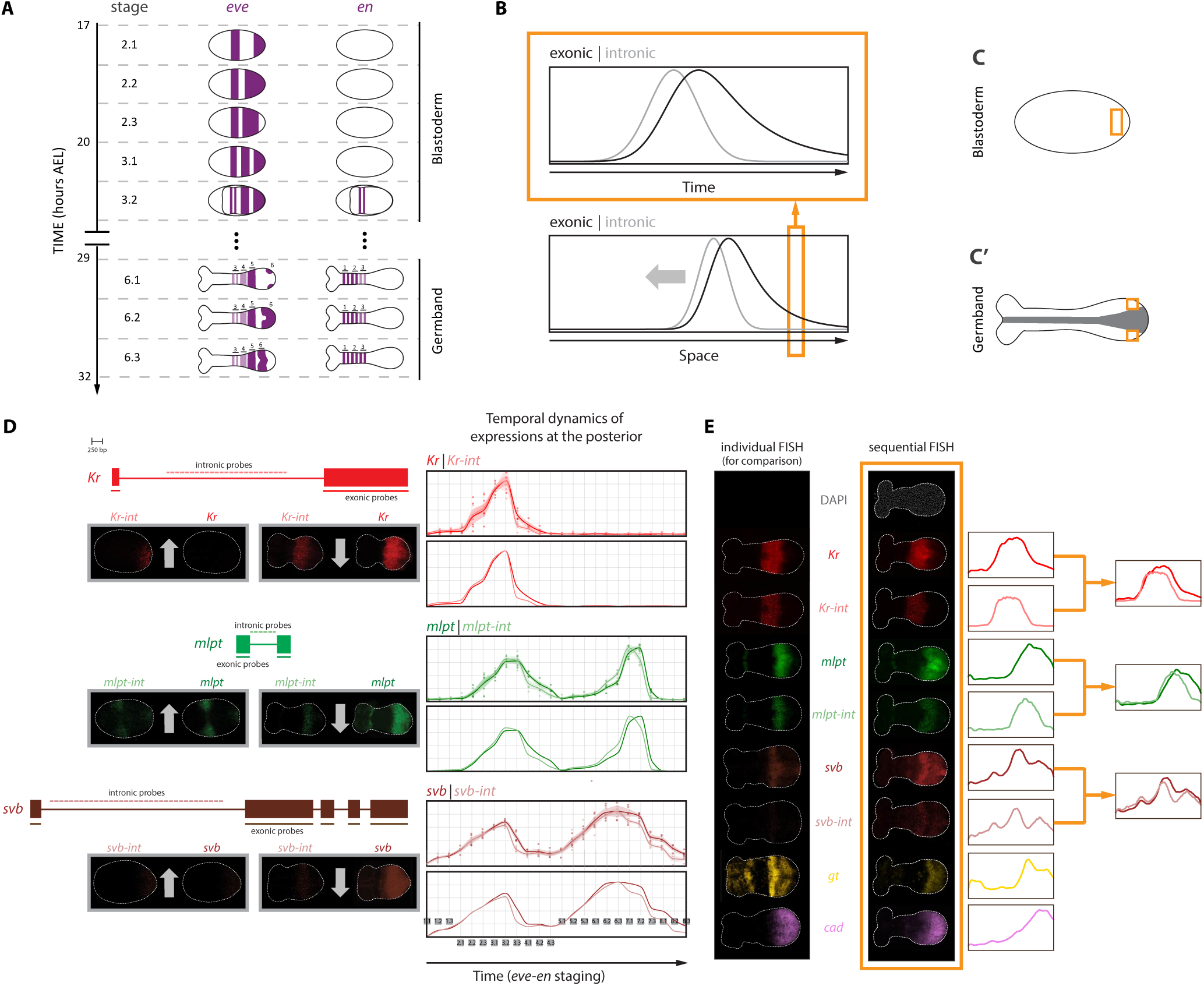
Intronic-exonic framework for inferring gene expression dynamics. (**A) Temporal staging.** Embryos were temporally aligned using the dynamic expression patterns of *eve* and *en*. *eve* is expressed in sequential oscillatory waves at the posterior signaling center, with each oscillatory cycle subdivided into three successive stages corresponding to the initial appearance, expansion, and posterior clearance of expression. Because each *eve* oscillatory cycle gives rise to a pair-rule stripe that subsequently splits into two segmental domains, counting eve cycles provides a temporal readout of developmental progression during early stages. At later stages, as mature eve stripes progressively resolve into segmental patterns and weaken anteriorly, stable en segmental stripes were used as complementary temporal landmarks. Together, eve oscillatory dynamics and en segmental domains establish a unified staging framework spanning both blastoderm and germband development. (**B) Simulation of propagating transcriptional signals.** We simulated a transcriptional pulse propagating through space with constant velocity and width (bottom panel), modeling intronic signal as a near-instantaneous readout of transcription and exonic signal as an accumulated transcript pool shaped by mRNA persistence and decay. The orange rectangle marks a fixed spatial position within the propagating domain, whose corresponding temporal profiles are shown in the top panel. Simulation shows that the temporal offset between intronic and exonic expression is smaller during the rising phase of the pulse and larger during the declining phase (top panel). As the pulse propagates through space, this temporal asymmetry is converted into a spatial phase shift between intronic and exonic signals (bottom panel), producing a smaller offset at the leading edge and a larger offset at the trailing edge of the propagating domain. (**C, C’) Quantification regions.** Gene expression levels were quantified within defined posterior regions corresponding to the posterior signaling center, where the AP temporal gene expression sequence is generated. In blastoderm embryos (C), measurements were obtained from a posterior region encompassing the site of sequential gene activation. In germband embryos (C’), quantification was restricted to the posterior ectoderm using two regions of interest to exclude mesodermal tissue, as mesodermal expression exhibits a slight temporal delay relative to the ectoderm. (**D) Validation of intronic-exonic offsets.** Left: Selected intronic and exonic regions of the gap genes *Kr*, *mlpt*, and *svb* used for HCR probe design are shown. Representative embryos stained simultaneously with intronic and exonic probes illustrate the relationship between nascent and accumulated transcripts during both activation (upward arrows) and deactivation (downward arrows) phases of gene expression. During activation, intronic expression appears before detectable exonic accumulation, whereas during deactivation, intronic expression clears from the posterior before exonic expression declines, consistent with the expected temporal relationship between nascent transcription and accumulated mRNA. Right: Quantification of intronic and exonic expression levels at the posterior end of the embryo across developmental stages for all analyzed gap genes. Upper plots show expression profiles together with individual data points (each dot representing one embryo) and confidence intervals. Intronic expression consistently precedes exonic expression during activation, although the temporal offset during the rising phase remains close to the noise level. In contrast, during the declining phase, exonic expression persists substantially longer than intronic expression, producing a larger and clearly detectable temporal offset. Nevertheless, the consistent early appearance of intronic signal across developmental stages supports the biological significance of the smaller offset during activation. Lower plots show the same expression profiles without scatter plots or confidence intervals for clarity of visualization. **(E) Gene expression spatiotemporal dynamics inferred through intronic-exonic comparison.** Comparison between conventional intronic-exonic HCR stainings and sequential multiplex imaging confirms that the sequential imaging protocol faithfully reproduces the intronic-exonic expression relationships observed using regular HCR stainings. Sequential imaging enables simultaneous visualization and direct spatial comparison of intronic and exonic expression domains for multiple gap genes within the same embryo. The observed asymmetric spatial offsets, characterized by smaller separation at the leading edge and larger separation at the trailing edge, match the computational predictions and indicate continued propagation of gene expression domains from posterior to anterior.

However, as *eve* stripes propagate anteriorly, they progressively broaden and weaken, limiting their utility for staging older embryos. To extend temporal staging into later developmental stages, we used *en* expression as a complementary marker. Each mature *eve* pair-rule stripe subsequently splits into two segmental domains that give rise to stable *en* stripes in the elongating germband. Counting *en* stripes therefore provides a reliable estimate of the number of completed *eve* oscillation cycles and enables temporal alignment of embryos across both blastoderm and germband stages (Fig. 3A). Together, this framework establishes a unified temporal coordinate system for analyzing AP patterning dynamics throughout development.

### 3. Intronic-exonic comparisons provide a framework for analyzing propagatory gene expression dynamics

#### 3.1 Intronic-exonic offsets as signatures of propagatory gene expression dynamics

Having established both a multiplex imaging framework for visualizing multiple AP patterning genes within the same embryo and a temporal staging system for aligning embryos across developmental progression, we next sought to extend this approach to capture the dynamics of these gene expression domains across space. Characterizing such dynamics is essential for understanding temporal-to-spatial patterning mechanisms, including the Speed Regulation model (Fig. 1C,C’), in which sequential temporal programs are transformed into spatial patterns through propagating waves of gene expression. Such propagatory behavior is a central feature of the gradient-based mode of the model, as well as tapered implementations of the wavefront-based mode discussed in the following section. To investigate these dynamics experimentally, we evaluated whether contrasting intronic and exonic signals could be used to infer gene expression behavior in the early *Tribolium* embryo. Because intronic signal reflects nascent transcription whereas exonic signal reflects accumulated mRNA, the temporal delay between them can be converted into a spatial offset when gene expression domains propagate through the tissue.

More generally, this strategy provides a readout of propagatory gene expression dynamics independent of the specific underlying model, and therefore offers a general framework for studying dynamic pattern formation in fixed embryos. To illustrate this behavior, we performed a computational simulation of a propagating transcriptional pulse with constant velocity and width, modeling the intronic signal as a near-real-time readout of transcriptional activity and the exonic signal as an accumulated transcript pool shaped by transcript processing and persistence (Fig. 3B). Because the exonic signal integrates recent transcriptional history, it does not follow the intronic signal with a uniform offset across the pulse. At a fixed spatial location, the exonic signal rises shortly after transcription initiates, resulting in a relatively small temporal offset at the leading edge of the pulse, but persists after transcription declines, producing a larger offset at the trailing edge (Fig. 3B, upper panel). As the transcriptional pulse propagates through space, this temporal asymmetry is converted into a spatial phase shift between intronic and exonic signals: at both edges of the pulse, the intronic signal leads the exonic signal, but the spatial offset between the two signals is relatively small at the leading edge and substantially larger at the trailing edge (Fig. 3B, lower panel). This asymmetric intronic-exonic relationship provides a characteristic signature of propagating gene expression domains and illustrates how intronic-exonic comparisons can be used to infer transcriptional dynamics across embryonic tissues.

#### 3.2 Measuring temporal intronic-exonic relationships for select AP patterning genes in *Tribolium*

To evaluate whether intronic-exonic comparisons can capture propagatory gene expression dynamics during AP patterning, we combined this approach with the temporal staging framework described above (Fig. 3A). We focused on three representative gap genes, *Kr*, *mlpt*, and *svb*, and designed HCR probe sets targeting both intronic and exonic regions of each gene (Fig. 3D). Embryos were co-stained with intronic and exonic probes using distinct fluorophores, enabling simultaneous visualization of nascent and accumulated transcripts within the same embryo (Fig. 3D).

Using the *eve-en* staging framework (Fig. 3A), embryos were temporally aligned across developmental progression, and intronic and exonic expression levels were quantified at the posterior (Fig. 3C,C’). For blastoderm stages, signal intensity was averaged within a defined posterior region (Fig. 3C). For germband stages, measurements were restricted to the posterior ectoderm using two regions of interest to exclude mesodermal contributions, as mesodermal expression exhibits a slight temporal delay relative to the ectoderm (Fig. 3C’). Background-subtracted intronic and exonic intensities were then plotted across developmental time (Fig. 3D).

For all genes analyzed (Fig. 3D), intronic expression preceded exonic expression during both activation and deactivation phases. At a fixed posterior location, intronic signal appeared earlier at the onset of expression and declined earlier as expression decreased, consistent with its role as a readout of nascent transcription (Fig. 3D). Importantly, the temporal offset between intronic and exonic signals was not uniform throughout the expression pulse. During the rising phase, intronic and exonic signals were closely aligned, whereas during the declining phase, the offset became substantially larger (Fig. 3D). This asymmetric behavior closely matches the predictions of our computational simulation (Fig. 3A), in which a propagating transcriptional pulse with a fixed temporal delay generates a smaller offset at the leading edge and a larger offset at the trailing edge.

Although the measured offset during the rising phase approached the noise level (Fig. 3D), its consistent appearance across developmental time supports its biological significance. In contrast, the larger offset during the declining phase was clearly distinguishable from the variance and readily detectable (Fig. 3D). Representative embryos illustrate these relationships directly, showing cases in which intronic signal appears before detectable exonic accumulation during activation, as well as cases in which exonic signal persists after intronic signal has diminished during deactivation (Fig. 3D).

Together, these results demonstrate that intronic-exonic comparisons provide a robust experimental readout of gene expression dynamics and validate their potential use for assessing the propagation of gene expression domains in the *Tribolium* embryo.

#### 3.3 Sequential multiplex imaging resolves propagatory dynamics within the spatial context of the embryo

Having established that intronic-exonic comparisons provide a reliable temporal readout at a fixed posterior location, we next applied our sequential imaging approach to examine these relationships in space within the same embryo (Fig. 3E). We simultaneously visualized a set of patterning genes, including both intronic and exonic expression of *Kr*, *mlpt*, and *svb*, together with exonic expression of *gt* and *cad* (Fig. 3E). This multiplexed dataset allowed direct comparison of multiple genes and their expression dynamics within the same embryo.

To validate the accuracy of the sequential imaging strategy, expression patterns obtained from multiplexed imaging were compared with conventional regular (not sequential) FISH stainings at corresponding developmental stages (Fig. 3E), confirming faithful reproduction of expected expression domains. Gene expression profiles were then quantified as described above, enabling systematic comparison of intronic and exonic signals across the AP axis.

Consistent with the temporal analysis, we observed a characteristic spatial offset between intronic and exonic expression domains. The separation between the two signals was smaller at the leading edge of each expression domain and larger at the trailing edge (Fig. 3E). This asymmetry matches the behavior predicted by our computational simulations and reflects the underlying temporal delay between transcriptional activation and mRNA accumulation (Fig. 3A). These results demonstrate that sequential imaging combined with intronic-exonic labeling can resolve gene expression dynamics within the spatial context of the embryo.

### 4. Intronic-exonic analysis reveals continued movement of gene expression domains during the germband stage

We next used our sequential FISH framework combined with intronic-exonic visualization to investigate the specific mode of temporal-to-spatial patterning in *Tribolium*. As described above, the Speed Regulation model provides a mechanistic framework for this process and has received strong experimental support for AP patterning in the early embryo. In this model, patterning can operate in two limiting modes (Fig. 1C,C’): a gradient-based mode, in which a non-retracting morphogen gradient modulates gene expression dynamics across a fixed tissue, and a wavefront-based mode, in which a retracting boundary stabilizes gene expression states in elongating tissues. In *Tribolium*, the candidate speed regulator *cad* is expressed as a non-retracting gradient in the blastoderm and as a retracting wavefront in the germband (Fig. 4A).

**Figure 4.**
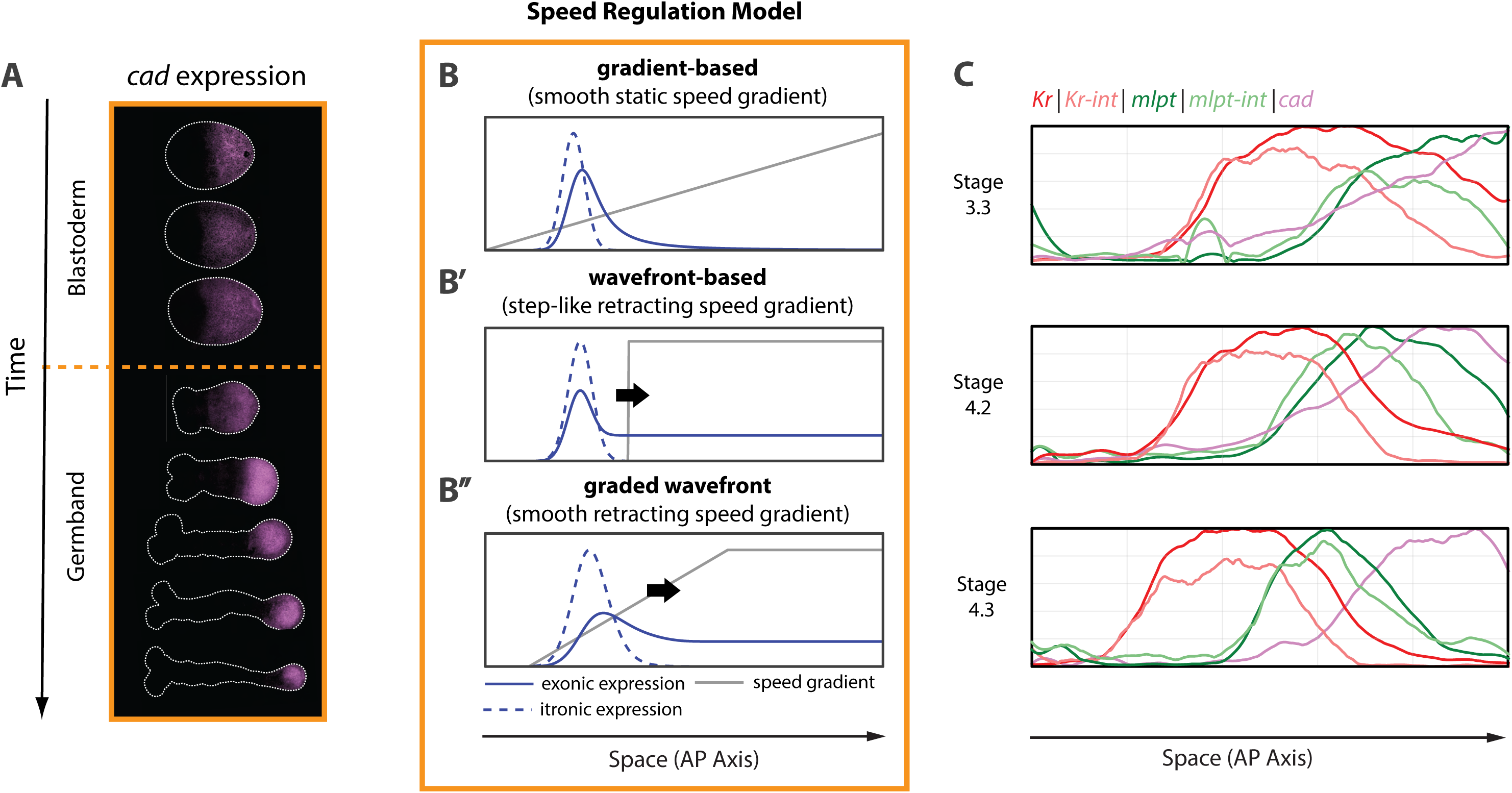
Continued propagation of gene expression domains in elongating tissues. **(A) Spatiotemporal expression of *cad***. The candidate speed regulator *cad* is expressed as a broad, non-retracting posterior-to-anterior gradient during blastoderm stages. During germband elongation, the *cad* domain progressively retracts posteriorly together with tissue elongation, transitioning into a wavefront-like configuration. **(B-B’’) Model predictions.** Simulations of the Speed Regulation model under three distinct regulatory regimes. In the static gradient regime (B), a non-retracting gradient continuously modulates progression through the temporal gene expression sequence, generating propagating waves of gene expression and clear intronic-exonic spatial offsets. In the sharp retracting wavefront regime (B’), cells anterior to the wavefront are rapidly stabilized as the wavefront retracts, resulting in little or no propagation and therefore minimal intronic-exonic spatial offset. In the graded retracting wavefront regime (B’’), propagation persists within the graded portion of the retracting boundary, producing detectable intronic-exonic spatial offsets despite overall wavefront retraction. Insets show the predicted intronic-exonic relationships for each regime.**(C) Experimental propagation in the germband.** Sequential intronic-exonic imaging of *Kr* and *mlpt* during germband elongation reveals consistent asymmetric spatial offsets across successive developmental stages. In both genes, intronic expression leads exonic expression, with smaller separation at the leading edge and larger separation at the trailing edge of the expression domains. The persistence of these offsets during tissue elongation indicates continued propagation of gene expression domains in the germband and is more consistent with a graded retracting wavefront than with a sharp stabilization boundary.

In addition to these two limiting cases, the model allows for an intermediate regime in which the retracting wavefront is graded rather than sharp. These different regimes make distinct predictions regarding gene expression dynamics (Fig. 4 B, B’, B’’). In the gradient-based case (Fig. 4B), gene expression domains propagate across the tissue. In the sharp wavefront case (Fig. 4B’), expression domains arise de novo as the wavefront retracts, with little or no propagation. In the intermediate case of a tapered (graded) wavefront (Fig. 4B’’), gene expression domains continue to propagate within the graded region of the retracting boundary. Because the key distinction between the latter two regimes lies in whether expression domains move, we asked whether intronic-exonic comparisons could discriminate between them.

To address this, we simulated the Speed Regulation model under all three scenarios (Fig. 4 B, B’, B’’). In the gradient-based regime, propagating gene expression domains produce a clear spatial offset between intronic and exonic signals (Fig. 4B). In the sharp wavefront regime, this offset is largely absent, reflecting the lack of domain propagation (Fig. 4B’). In contrast, the tapered wavefront regime retains a spatial offset, consistent with continued propagation within the graded region (Fig. 4B’’). These results indicate that intronic-exonic comparisons provide a means to distinguish between sharp and graded wavefront implementations of the model.

In previous work, we showed that gap gene expression domains propagate across the blastoderm. This was possible to establish because the blastoderm is a non-elongating tissue, allowing spatial positions of gene expression domains to be compared across embryos. In the germband, however, continuous tissue elongation complicates such analysis, as apparent movement of expression domains can arise from both gene expression dynamics and tissue deformation. This has limited direct assessment of propagation in elongating tissues.

Using the intronic-exonic framework, we next examined the dynamics of selected gap genes during the *Tribolium* germband stage. For both *Kr* and *mlpt*, we observed consistent spatial offsets between intronic and exonic expression domains, indicating that these genes continue to propagate during germband elongation (Fig. 4C). Notably, intronic-exonic shifts for both genes were detected within the same posterior region, particularly within the domain of detectable *cad* expression (Fig. 4C), indicating that multiple gap genes concurrently propagate away from the posterior signaling center. Because multiple genes propagate simultaneously within the same developmental and spatial context, this raises the possibility that differences in effective propagation dynamics between genes could contribute to changes in relative spatial phasing as expression domains move away from the posterior, a possibility discussed further below.

Together, these observations demonstrate that gap gene expression domains continue to exhibit propagatory behavior during the germband stages of *Tribolium* development. Thus, temporal-to-spatial patterning in the AP system remains dynamically active throughout tissue elongation rather than becoming spatially fixed once initial domains are established.

### 5. Assessing the fidelity of temporal-to-spatial transformation in patterning

Temporal-to-spatial patterning entails the conversion of a temporal gene expression program into a spatial pattern along the developing tissue. In its simplest realization, this transformation would be faithful, such that the temporal relationships between genes are preserved in space, resulting in corresponding spatial relationships among their expression domains. However, whether temporal information is directly translated into spatial organization or progressively reshaped during propagation remains unclear. Having established a sequential imaging strategy capable of simultaneously analyzing multiple genes together, we next sought to directly compare temporal gene activation at the posterior with the resulting spatial organization of patterning genes along the AP axis.

#### 5.1 Temporal profiling of gap and pair-rule genes at the posterior

To address this question, it was first necessary to characterize the temporal sequence of gene activation at the posterior of the *Tribolium* embryo, where patterning dynamics originate, and then compare this sequence to the spatial arrangement of gene expression domains. We focused on primary pair-rule genes (*eve*, *run*, and *odd*) and gap genes (*hb*, *kr*, *mlpt*, *gt*, and *svb*).

To establish the temporal sequence, we quantified gene expression levels at the posterior using conventional FISH (Fig. 5A). For each gene, measurements were obtained from multiple embryos per developmental stage (n = 5), using the same quantification framework described above (Fig. 3C). Embryos were staged using the *eve*-*en* system (Fig. 3A), allowing alignment across developmental time. This analysis yielded temporal expression profiles for both pair-rule and gap genes at the posterior of the embryo (Fig. 5B).

**Figure 5.**
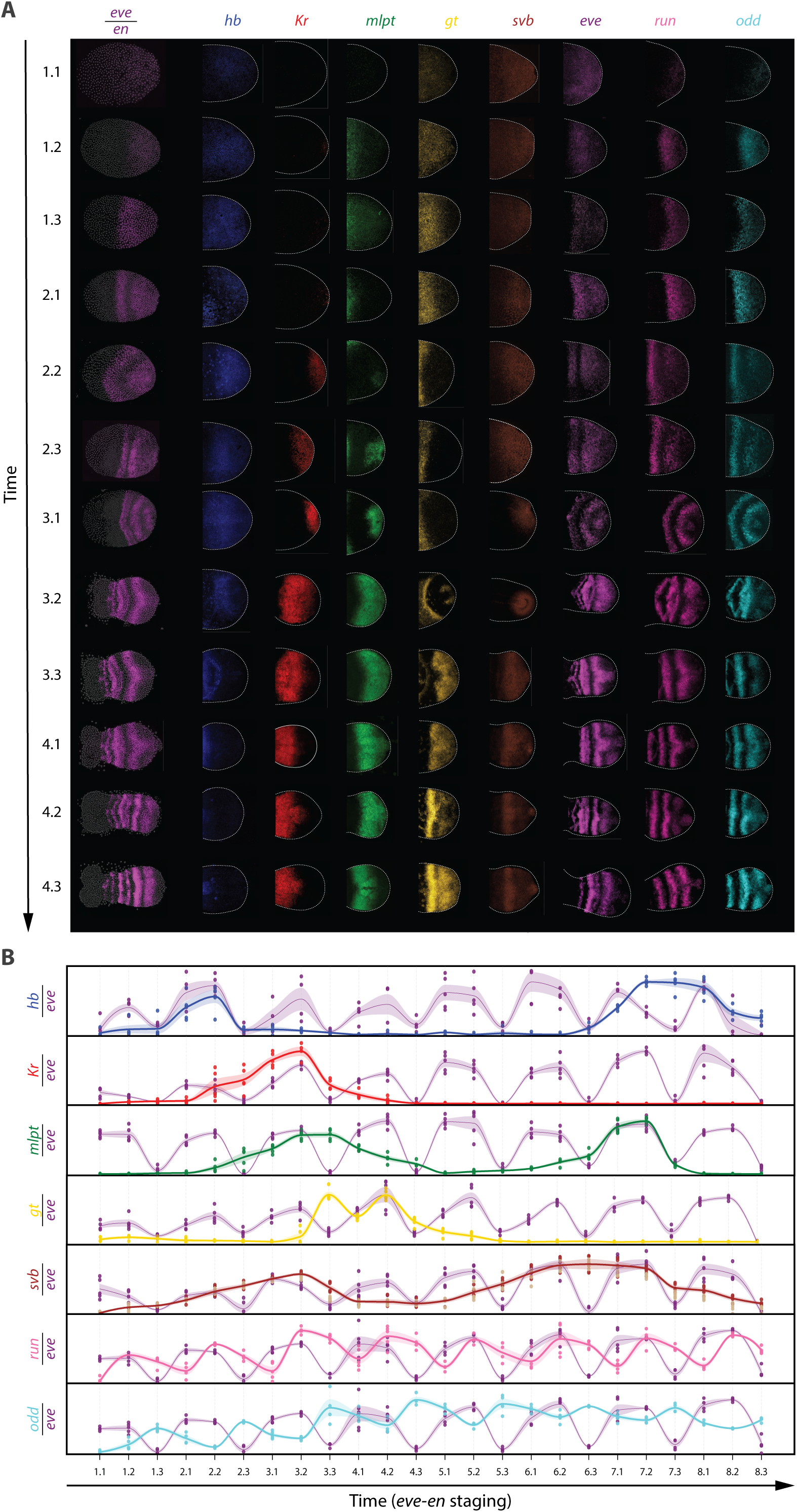
Quantitative temporal profiling of AP patterning genes at the posterior. **(A) Representative temporal gene expression sequence**. Representative embryos staged using the eve-en temporal framework are shown to illustrate the sequential activation dynamics of gap and pair-rule genes at the posterior signaling center. Pair-rule genes exhibit cyclic oscillatory expression, whereas gap genes display sequential but non-periodic activation dynamics. Together, these expression patterns define the temporal progression of AP patterning at the posterior of the embryo. **(B) Quantitative temporal profiles.** Background-subtracted expression intensities measured at the posterior signaling center were quantified across developmental stages defined by the eve-en staging system. Temporal profiles reveal the onset, peak, duration, and decline of expression for each gene, establishing the relative temporal ordering and phase relationships between patterning genes during AP patterning. These profiles provide the temporal reference framework used for subsequent comparison between posterior gene activation dynamics and the resulting spatial organization of expression domains along the AP axis.

#### 5.2 Assaying the spatial relationships of patterning genes across developmental stages

To relate the temporal sequences described above to spatial gene expression patterns, we applied our sequential FISH approach to *Tribolium* embryos across successive developmental stages, defined using the *eve*-*en* staging scheme (stages 3.3, 4.1, 4.3, and 5.2) (Fig. 6). We simultaneously visualized a panel of patterning genes (*hb, Kr, mlpt, gt, svb, eve, run, odd, en*, and *cad*), enabling direct assessment of their spatial relationships and how these relationships evolve over time (Fig. 2C, Fig. 6).

**Figure 6.**
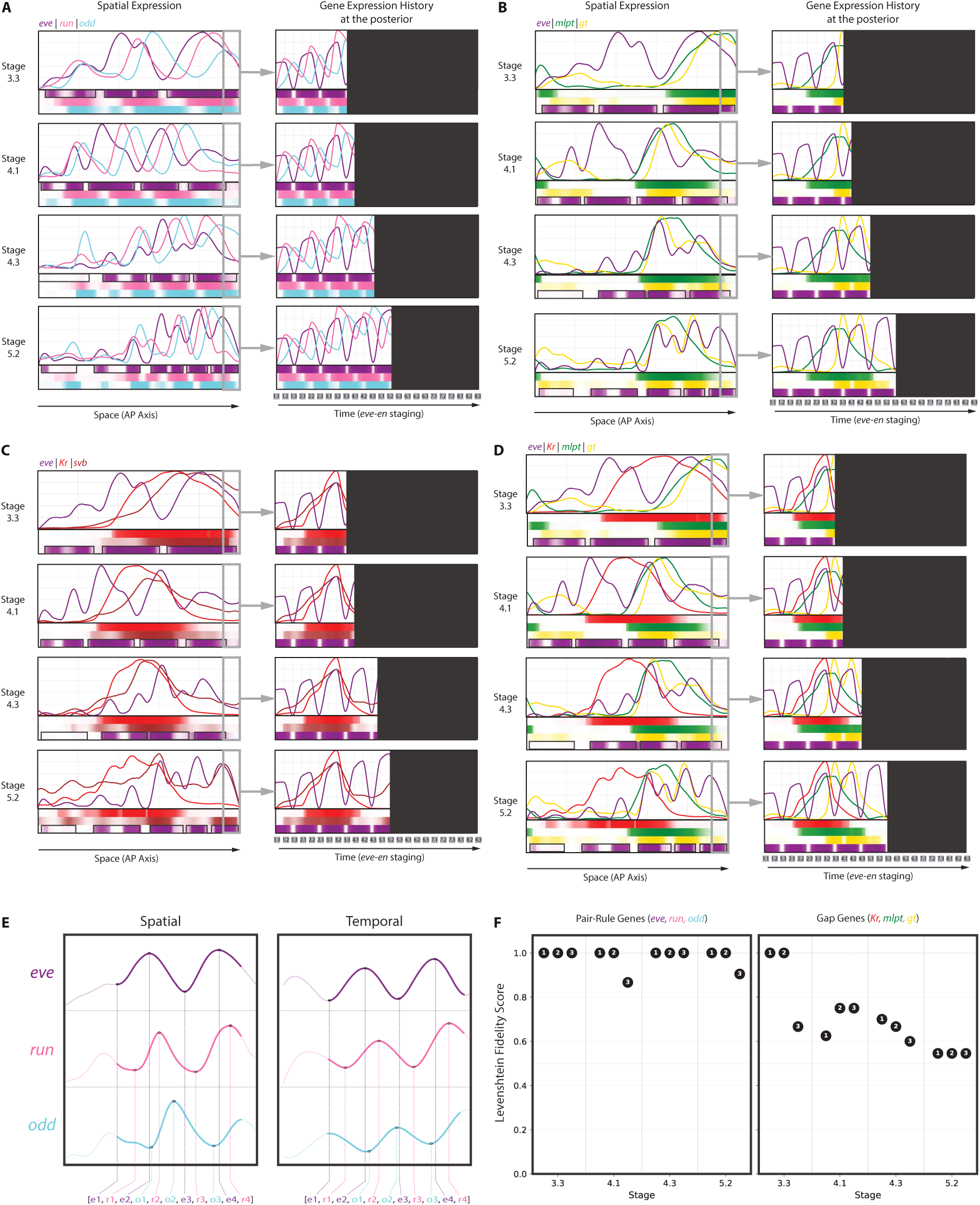
Comparison between temporal activation of AP patterning genes at the posterior and their resulting spatial organization along the AP axis. Right panels show the temporal sequence of AP patterning gene activation at the posterior signaling center, reconstructed from the quantitative temporal profiles in Fig. 5. Left panels show the corresponding spatial organization of gene expression domains along the AP axis, resolved using sequential multiplex imaging at developmental stages 3.3, 4.1, 4.3, and 5.2 according to the eve-en staging framework. For each stage, temporal plots include only the gene expression history that occurred up to the indicated developmental stage. Block representations below each panel summarize the corresponding temporal and spatial phase relationships between genes (see Materials and Methods for generation of block representations). **(A) Faithful temporal-to-spatial transformation in primary pair-rule genes.** The temporal sequence of *eve*, *run*, and *odd* oscillatory pulses at the posterior signaling center (right) is preserved in their spatial arrangement along the AP axis (left), indicating that relative phase relationships are maintained during spatial pattern formation. **(B-D) Non-faithful temporal-to-spatial transformation in gap genes.** Temporal activation profiles of selected gap genes are compared with their resulting spatial organization relative to *eve* expression. (B) Comparison of *eve*, *mlpt*, and *gt*. (C) Comparison of *eve*, *Kr*, and *svb*. (D) Comparison of *eve*, *Kr*, *mlpt*, and *gt*. In contrast to pair-rule genes, the relative spatial relationships between gap genes progressively diverge from their temporal relationships established at the posterior signaling center. **(E) Landmark extraction and sequence representation.** Temporal and spatial expression patterns were converted into ordered sequences of landmark events to enable quantitative comparison. Peaks and troughs were identified from temporal and spatial profiles of the primary pair-rule genes *eve*, *run*, and *odd* and represented as ordered event lists. The resulting sequences preserve the relative ordering of prominent expression landmarks while remaining insensitive to differences in absolute spacing between domains, allowing direct comparison between temporal and spatial organization. **(F) Quantification of temporal-to-spatial faithfulness.** Temporal and spatial event sequences were compared using a faithfulness metric based on normalized Levenshtein distance. Faithfulness scores were calculated for embryos at stages 3.3, 4.1, 4.3, and 5.2 (three biological replicates per stage), where a score of 1 indicates perfect preservation of event ordering and lower values indicate increasing divergence between temporal and spatial organization. Pair-rule genes maintain high faithfulness scores across developmental stages, whereas gap genes exhibit progressively reduced faithfulness, quantitatively confirming the differential temporal-to-spatial transformation revealed by the qualitative analyses.

This dataset reveals several notable features. For clarity, we first examine selected subsets of genes separately before integrating these observations into an overall view of the system-level behavior.

#### 5.3 Faithful translation of temporal sequence into spatial pattern in pair-rule genes

We first examined the temporal relationships between the primary pair-rule genes in *Tribolium* (*eve, run,* and *odd*; Fig. 6A, right panel) and compared them to their corresponding spatial organization (Fig. 6A, left panel). At the posterior, pair-rule gene expression is oscillatory, producing a series of eight sequential pulses that give rise to the eight primary pair-rule stripes along the AP axis. These pulses exhibit a consistent temporal phase relationship: each cycle is initiated by *eve*, followed by *run*, and then *odd* (Fig. 6A, right panel).

We next asked how these temporal relationships are reflected in space. The three primary pair-rule genes form periodic spatial patterns along the AP axis (Fig. 6A, left panel). Near the posterior, where new expression domains emerge, the spatial arrangement mirrors the temporal sequence: within each repeating unit, domains are ordered from anterior to posterior as *eve*, followed by *run*, and then *odd*. As these domains propagate anteriorly, they undergo refinement: *eve* stripes split into two segmental domains, while *odd* resolves into segmental pattern, and all domains eventually fade. Despite these transformations, the relative phase relationships established at the posterior are largely preserved as gene expression waves emanate from the posterior (compare right and left panels in Fig. 6A), indicating that the temporal sequence of pair-rule gene activation is faithfully translated into spatial organization.

#### 5.4 Gap genes exhibit non-faithful temporal-to-spatial transformation

We next examined gap genes to determine whether their temporal activation is similarly reflected in their spatial organization. As above, we compared temporal expression dynamics at the posterior (Fig. 6B-D; right panels) with spatial expression patterns along the AP axis (Fig. 6B-D; left panels), focusing on subsets of genes one at a time for clarity.

We first analyzed the temporal relationship between *mlpt* and *gt* (Fig. 6B, right panel). At the posterior, *mlpt* expression is initiated early (around stage 2.2) and persists until approximately stage 4.3, after which it is cleared. A second phase of *mlpt* expression appears at later stages (Fig. 6B right panel). In contrast, *gt* expression is initiated later (around stage 3.2), following *mlpt*, and exhibits two successive pulses, with the second pulse extending beyond the temporal window of *mlpt*. Thus, temporally, *mlpt* precedes *gt*, and their expression windows partially overlap, with *gt* extending later in time.

We then compared these temporal relationships to their spatial organization (Fig. 6B, left panel). Contrary to the expectation of a faithful temporal-to-spatial mapping, the spatial arrangement does not directly reflect this temporal sequence. At stage 4.3, where both genes are active, the second phase of *gt* does not extend posteriorly beyond the *mlpt* domain as predicted from its temporal profile. Instead, *gt* is positioned within or anterior to the *mlpt* domain. More generally, *mlpt* appears spatially restricted toward posterior regions relative to its temporal profile, whereas *gt* is positioned more anteriorly than expected. These observations indicate that the temporal relationships between *mlpt* and *gt* are progressively reshaped during the conversion from temporal sequence into spatial pattern.

We next examined the relationship between *Kr* and *svb* (Fig. 6C). Temporally (Fig. 6C, right panel), *svb* expression is initiated slightly earlier than *Kr* (around stages 1.2-2.1) and persists slightly longer (until approximately stage 4.1). Despite this, their peak expression levels largely coincide around stage 3.2. However, this temporal relationship is not preserved in space. Spatially, *svb* is positioned more posteriorly than *Kr* (Fig. 6C, left panel), indicating that the relative phase relationship between the two genes changes during the conversion of the temporal sequence into spatial organization.

Comparison of multiple gap gene expression patterns together with *eve* further supports these observations (Fig. 6D). Temporally, *Kr* expression overlaps with the mid-second cycle of *eve*, followed shortly by *mlpt*, whereas *gt* expression begins later, overlapping with late cycle 3 of *eve*. However, these temporal relationships are not preserved in space (Fig. 6D, stage 4.3). Spatially, *Kr* overlaps with the onset of *eve* cycle 2, while *mlpt* aligns with the onset of cycle 3, indicating that *mlpt* is positioned more posteriorly than expected relative to *Kr*. A similar discrepancy is observed for *gt*. Temporally, the first *gt* pulse occurs between *eve* cycles 3 and 4, while the second pulse overlaps with cycles 4 and 5. In contrast, spatially, the first *gt* domain aligns with the third *eve* stripe, and the second domain aligns with the fourth stripe. Thus, relative to its temporal activation profile, *gt* occupies more anterior positions than expected. Together, these comparisons show that the relative phase relationships between gap genes and *eve* are progressively altered during the conversion of temporal sequence into spatial organization.

#### 5.5 Quantitative verification of faithfulness of temporal-to-spatial transformation

Although the analyses above reveal clear qualitative differences between pair-rule and gap genes, quantifying temporal-to-spatial transformation is not straightforward because temporal and spatial patterns differ in multiple respects, many of which are not directly relevant to pattern identity. Since the biological outcome of pattern formation is the establishment of correctly ordered spatial domains, we focused on the relative ordering of prominent expression landmarks (specifically, peaks and troughs) across gene expression profiles (both spatial and temporal).

We first identified these landmarks in both temporal and spatial profiles and represented them as ordered sequences of events (see example in Fig. 6E). The resulting temporal and spatial sequences were then compared using a faithfulness metric based on normalized Levenshtein distance, which quantifies the similarity of ordered event sequences (see Methods). A score of 1 indicates perfect preservation of event ordering, whereas lower scores indicate progressive divergence between temporal and spatial organization.

We applied this analysis to pair-rule and gap gene datasets obtained by sequential imaging at stages 3.3, 4.1, 4.3, and 5.2, using three biological replicates per stage. Pair-rule genes consistently exhibited high faithfulness scores across developmental stages, indicating preservation of temporal phase relationships during spatial pattern formation (Fig. 6F). In contrast, gap genes displayed substantially lower faithfulness scores, with divergence becoming increasingly pronounced at later stages. These results quantitatively support the conclusion that pair-rule genes undergo largely faithful temporal-to-spatial transformation, whereas gap gene patterns are progressively reshaped during propagation.

One possible interpretation of these observations is that different gap genes exhibit distinct effective propagation dynamics as their expression domains move from posterior to anterior. Consistent with our earlier intronic-exonic analysis demonstrating continued propagation of gap gene expression domains across developmental stages, this interpretation would imply that some gene expression domains advance anteriorly more rapidly relative to their temporal activation profiles, whereas others remain spatially confined to more posterior regions. As a result, the relative spatial phasing between genes would progressively diverge from the temporal relationships originally established at the posterior signaling center.

However, because these genes operate within a shared GRN, such differences are unlikely to arise solely from independent gene-specific behaviors. Instead, the observed reshaping of spatial phase relationships may reflect progressive changes in the underlying regulatory interactions as patterning proceeds from posterior to anterior. In this view, temporal-to-spatial patterning would involve not only propagation of gene expression domains, but also dynamic reconfiguration of GRN structure during development. We explore these possible mechanisms further in the Discussion.

## Discussion

To investigate how temporal gene expression programs are transformed into spatial patterns, we developed a sequential multiplexed imaging strategy based on HCR, enabling visualization of up to ten genes within the same *Tribolium* embryo (Fig. 2). By combining this approach with intronic-exonic labeling (Fig. 3), we established a framework for inferring gene expression dynamics and assessing propagatory behavior of expression domains in fixed embryos. This framework further enables direct comparison between temporal gene activation at the posterior signaling center (Fig. 5) and the resulting spatial organization of gene expression domains along the AP axis (Fig. 6).

We first used intronic-exonic comparisons (Fig. 3) to directly assess gene expression dynamics (Fig. 4). Consistent with previous observations in the blastoderm, we find that gap gene expression domains continue to propagate in the germband, along with ongoing tissue elongation. This establishes that spatial patterns are dynamically reshaped during development rather than being passively fixed.

We then characterized the temporal sequence of gap and pair-rule gene activation at the posterior (Fig. 5) and compared it to their spatial arrangement (Fig. 6). We showed that primary pair-rule genes exhibit a faithful transformation from temporal sequence to spatial pattern, preserving phase relationships as expression domains propagate (Fig. 6A). In contrast, gap genes display systematic deviations from this mapping: their spatial domains undergo gene-specific changes in relative positioning compared to the temporal relationships established during posterior activation (Fig. 6B-D).

Together, these findings indicate that temporal-to-spatial patterning is not uniformly implemented across the different AP patterning systems analyzed here. While pair-rule genes preserve temporal relationships in space, gap genes undergo active transformation during propagation, resulting in altered spatial phasing. This raises the question of what mechanisms underlie these gene-specific differences and how they contribute to the establishment of robust spatial patterns during development.

### Possible mechanisms

A natural starting point for explaining this behavior is the Speed Regulation model, in which a morphogen gradient modulates the timing of a temporal gene sequence and thereby generates propagating spatial patterns. One direct implementation of such timing control at the level of a GRN is through coordinated modulation of transcription rates and gene product decay rates across all constituent genes, effectively scaling the speed of the entire network, as shown previously [24]. Extending this idea, one possible explanation for the observed phase shifts is that different genes interpret the speed-regulating gradient differently, resulting in gene-specific effective speed profiles. To explore this possibility, we generated simulations in which temporal gene expression pulses were regulated by gradients of different shapes. These simulations showed that more concave gradient profiles can produce faster propagation, causing some spatial domains to advance ahead of others (Fig. 7A). This suggests that differential interpretation of a common upstream gradient could, in principle, generate the observed differences in spatial phasing.

**Figure 7.**
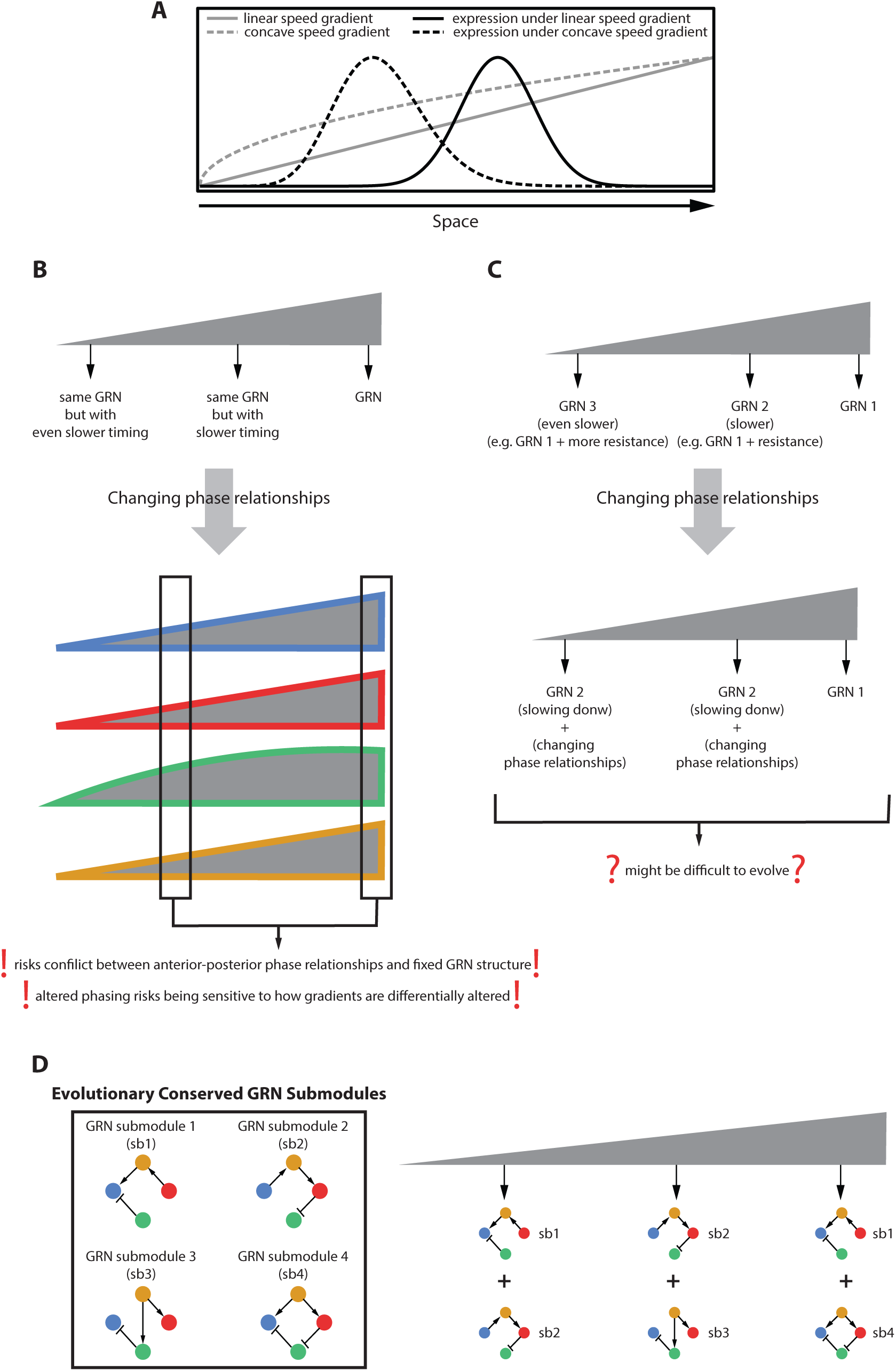
Mechanistic models for non-faithful temporal-to-spatial transformation. (A) Differential interpretation of speed gradients. Simulations of propagating temporal gene expression pulses regulated by gradients of different shapes show that more concave gradients generate faster anterior progression compared to linear gradients. As a result, gene expression domains regulated by distinct effective gradient profiles can become spatially shifted relative to one another despite originating from the same posterior temporal sequence, providing one possible explanation for the observed reshaping of spatial phase relationships. **(B) Limitations of independent temporal regulation of genes within a single GRN.** Schematic illustration of the challenges associated with explaining non-faithful temporal-to-spatial transformation solely through independent gene-specific propagation dynamics. In the hypothetical example shown, blue, red, and orange genes are regulated by linear effective speed gradients, whereas the green gene is regulated by a more concave gradient profile, resulting in faster propagation. While such differences could in principle generate altered spatial phase relationships, implementing distinct effective speed gradients for individual genes within a shared GRN risks destabilizing coordinated regulatory interactions and generating highly sensitive or unpredictable patterning outputs. **(C) GRN switching model. Schematic representation of progressive changes in regulatory interactions along the AP axis.** In this framework, cells initially operate within a dynamic posterior regulatory regime associated with sequential temporal progression, but progressively transition into distinct anterior regulatory configurations during propagation. Such GRN reconfiguration can alter relative timing and phase relationships between genes during spatial pattern formation, thereby reshaping the final spatial organization relative to the original temporal sequence generated at the posterior signaling center. **(D) Evolution and spatiotemporal modularity.** Conceptual framework illustrating how evolutionary selection may primarily act on the final spatial pattern rather than on faithful preservation of the initial temporal sequence itself. In this view, generating the required final spatial arrangement directly from a single posterior temporal GRN may be difficult because a single regulatory configuration may not readily produce the full set of required spatial phase relationships. Instead, spatial pattern formation may proceed progressively through sequential deployment and exchange of evolutionarily conserved GRN submodules along the AP axis. Posterior patterning dynamics would initially emerge from one combination of conserved regulatory submodules, which are then progressively replaced or reconfigured during anterior progression to generate the final spatial gene expression pattern. This temporal modularity framework provides a potential explanation for how non-faithful temporal-to-spatial transformation can arise through progressive rewiring of regulatory interactions during development.

However, this mechanism has important limitations. First, simulations in Fig. 7A consider isolated pulses, whereas gap genes operate within a shared GRN. Imposing gene-specific propagation speeds could therefore perturb network interactions, particularly in regions of low morphogen concentration, potentially leading to unpredictable outputs that depend on the underlying GRN wiring (Fig. 7B). Second, such a mechanism would be highly sensitive to the precise way each gene interprets the morphogen gradient, making it difficult to achieve the robust and precise spatial phasing required for downstream processes such as Hox gene regulation.

An alternative explanation is provided by the GRN switching model [16], [24]. In this framework, patterning is controlled by a progressive transition between distinct regulatory regimes along the AP axis: a dynamic GRN operating in the posterior drives sequential gene activation, while a multi-stable GRN in the anterior stabilizes spatial domains. The transition between these regimes is controlled by the same speed regulator gradient (Fig. 7C). In its original formulation, this model accounts for the progressive slowing and stabilization of gene expression dynamics [24]. A natural extension is that, as GRN wiring is progressively altered along the axis, not only timing but also phase relationships between genes could change, thereby reshaping spatial patterns during propagation (Fig. 7C).

At first glance, such a mechanism may appear difficult to evolve, as it would require GRN configurations capable of producing controlled changes in phase relationships during pattern formation (Fig. 7C). However, this difficulty assumes that the posterior temporal GRN is fixed and that subsequent regulatory modifications must progressively build upon it. A more plausible scenario is that evolutionary selection primarily acts on the final spatial pattern itself, while both posterior and anterior GRN configurations remain evolutionarily flexible (Fig. 7D).

This perspective raises a broader question: why does the initial temporal program not simply encode the final spatial pattern directly? One possibility is that GRNs possess only a limited repertoire of functional regulatory configurations, constraining the range of spatial relationships that can be generated by a single temporal GRN operating at the posterior (Fig. 7D). Under such constraints, the system may instead progressively transform the initial temporal output as gene expression domains move across the tissue. In this view, gap genes may operate through a limited set of reusable regulatory submodules (Fig. 7D), whose combinations are deployed differently along the AP axis under the influence of the speed regulator. Progressive changes in GRN wiring would therefore not only slow or stabilize gene expression dynamics, but also reshape phase relationships between genes during pattern formation. This would effectively implement a form of spatiotemporal modularity, in which the same genes participate in different regulatory configurations over time and space. Consistent with this idea, previous work has shown that regulatory interactions among gap genes in *Tribolium* differ between posterior and anterior regions [21], [58], [59]. Furthermore, it has been proposed that temporal-to-spatial patterning of insect gap genes evolved from ancestral temporal patterning programs operating in insect neuroblasts, where several gap genes are likewise activated sequentially and, strikingly, in an order corresponding to their temporal and spatial arrangement during AP patterning [15], [60]. However, the AP patterning system appears to have undergone substantial evolutionary modification through incorporation of additional genes into the sequence, such as *mlpt* and *gt*. These observations support the idea that temporal-to-spatial patterning systems may evolve through progressive reorganization and recombination of conserved regulatory interactions rather than through preservation of a fixed ancestral temporal program. However, this remains a working hypothesis. Future studies comparing the GRN architectures underlying gap gene regulation across space and time during both AP patterning and neurogenesis in *Tribolium* and other insects will be required to test this idea. Such analyses may provide important insights into the flexibility, modularity, and evolvability of developmental GRNs during embryogenesis.

Interestingly, during vertebrate somitogenesis, the phase relationship between oscillatory signaling activities progressively changes along the AP axis, and experimental evidence suggests that developmental information is encoded in these shifting phase relationships [61]. This raises the possibility that the progressive changes in phase relationships observed between *Tribolium* gap gene expression dynamics may likewise be genetically encoded and functionally important, rather than representing passive distortions arising during propagation.

Together, these considerations suggest that the observed non-faithful temporal-to-spatial transformation may either represent an evolutionary byproduct of the regulatory steps required to generate the final spatial pattern, or itself constitute a functionally important feature of pattern formation, as suggested by the progressive reshaping of phase relationships during vertebrate somitogenesis. The mechanisms underlying these changing phase relationships remain unclear. They may arise from differential effective propagation dynamics between genes, from progressive reconfiguration of the underlying GRN during pattern formation, or from a combination of both mechanisms. Additional mechanisms may also contribute. More broadly, however, our results indicate that temporal-to-spatial patterning can involve active transformation of temporal information rather than a simple direct mapping, revealing an additional regulatory layer underlying embryonic pattern formation.

## Movie legends

**Movie S1. Travelling transcriptional pulse along the anterior-posterior axis and its temporal readout at a fixed posterior location.** Shown is a simulation (from which the snapshots in Fig. 3B are taken) of a Gaussian intronic pulse travelling at constant speed from the posterior to the anterior end of the normalised AP axis. The exonic signal integrates transcriptional activity following first-order mRNA kinetics. The exonic profile is rescaled at each frame to match the peak amplitude of the intronic signal for visual comparison. Intronic and exonic signals are shown in gray and black, respectively.

**Movie S2. Speed regulation under a gradient-based morphogen.** Shown is a simulation (from which the first snapshot in Fig. 4B is taken) of a single-gene speed regulation system in which the morphogen is a static linear gradient, constant in time, increasing from zero at the anterior to its maximum at the posterior. The intronic and exonic signals are shown in blue. The morphogen gradient is shown in gray.

**Movie S3. Speed regulation under a wavefront-based morphogen.** Shown is a simulation (from which the second snapshot in Fig. 4B is taken) of a single-gene speed regulation system in which the morphogen is a sharp step function retracting posteriorly at constant speed. Cells to the right of the boundary receive full morphogen exposure; cells to the left receive none. The intronic and exonic signals are shown in blue. The morphogen gradient is shown in gray.

**Movie S4. Speed regulation under a graded wavefront.** Shown is a simulation (from which the third snapshot in Fig. 4B is taken) of a single-gene speed regulation system in which the morphogen is a smooth step function whose boundary retracts posteriorly over time. The intronic and exonic signals are shown in blue. The morphogen gradient is shown in gray.

**Movie S5. Effect of gradient shape on gene expression domain widths along the anterior-posterior axis.** Shown is a simulation (from which the snapshot in Fig. 7A is taken) of a single-gene speed regulation system comparing a static linear gradient and a static concave power-law gradient simultaneously. Linear gradient and its resulting expression pulse are shown as solid gray and solid black lines, respectively. Concave gradient and its resulting expression pulse are shown as dashed gray and dashed black lines, respectively.

## Materials and Methods

### Beetle cultures

Beetle cultures (GA-2 strain) were reared on flour supplemented with 5% dried yeast in a temperature– and humidity-controlled room at 24°C. To speed up development, beetles were reared at 32°C.

### Fixation of embryos

*Tribolium castaneum* eggs were collected, floated off in 50% bleach (∼2 min), rinsed in water, and fixed at room temperature for 1 h in a two-phase formaldehyde/heptane mixture. Devitellinization was achieved by rapid shaking with methanol, followed by repeated passage through a 20-gauge needle. Embryos were washed three times in methanol and stored at −20°C.

### HCR FISH

Embryos stored in methanol were rehydrated through 50% MeOH/PBT into PBT (1× PBS, 0.1% Tween-20), post-fixed in 4% formaldehyde/PBT for 30 min and rinsed in PBT. Embryos were pre-hybridized for 30 min in 30% probe hybridization buffer at 37°C, then incubated overnight with gene-specific HCR probes (Molecular Instruments) at 37°C. After four 15 min washes in 30% probe wash buffer (37°C) and three 5 min washes in 5× SSCT, snap-cooled fluorescent hairpins were applied in amplification buffer and allowed to react overnight at room temperature in the dark. Excess hairpins were removed by sequential washes in 5× SSCT, and embryos were mounted in SlowFade™ Gold (Thermo Fisher Scientific).

### Confocal Z-stacks were acquired using a Nikon AX confocal microscope

Images were collected with a field of view of approximately 686 × 448 µm at a lateral sampling resolution of 0.49 µm/pixel (2.01 pixels/µm in X and 2.02 pixels/µm in Y). Z-stacks consisted of approximately 42 optical sections acquired at a spacing of 0.65 µm. Maximum-intensity projections were generated in Fiji/ImageJ for visualization and downstream analysis.

### High multiplexity sequential imaging

Sequential imaging was performed using a modified hybridization chain reaction fluorescence in situ hybridization (HCR FISH) protocol adapted to enable repeated rounds of imaging within the same *Tribolium castaneum* embryo. Embryos were collected, fixed, devitellinized, and hybridized according to the standard procedures described above. Probe sets targeting multiple genes were designed using orthogonal HCR initiator barcode systems (B1-B10), and all probe sets were hybridized simultaneously prior to embryo mounting and imaging. Following hybridization, embryos were dissected and transferred onto PBS agarose gel pads. Chrome alum gelatin adhesive-coated circular coverslips were then gently pressed against the embryos on the agarose pad, allowing embryos to adhere to the coverslip surface while preserving their orientation. Stable immobilization of embryos was essential to prevent sample loss during repeated fluid exchange within the imaging chamber and to preserve embryo morphology throughout multiple rounds of amplification, imaging, bleaching, and reagent exchange.

Coverslips containing mounted embryos were subsequently assembled into a Bioptechs FCS2 imaging chamber, enabling controlled reagent exchange while maintaining embryo positioning and morphology throughout the sequential imaging procedure.

Before each amplification step, embryos were incubated in HCR amplification buffer. Fluid exchange within the Bioptechs FCS2 imaging chamber was performed manually by pipetting solutions through the chamber inflow inlet, while excess liquid exited through the outlet port. Fluorescent HCR hairpin amplifiers complementary to selected barcode initiators were then introduced in subsets corresponding to individual imaging rounds. In each round, up to three genes were visualized simultaneously using spectrally distinct fluorophore-conjugated hairpins, including Alexa Fluor 488, Alexa Fluor 546, and Alexa Fluor 647. Hairpins were snap-cooled immediately prior to use according to standard HCR procedures to ensure proper folding and amplification efficiency. After amplification, embryos were washed extensively through repeated buffer exchange within the chamber and imaged using confocal microscopy under identical acquisition settings within experimental series whenever possible.

Following imaging, fluorescent signal was removed through a combination of chemical bleaching and confocal photobleaching. Embryos were first subjected to chemical bleaching (see details of chemical bleaching below) to remove the majority of fluorophore signal. Residual fluorescence was subsequently eliminated directly on the confocal microscope using repeated high-intensity laser scanning for all imaging channels until no detectable signal remained. This combined bleaching strategy eliminates signal carryover between sequential imaging rounds. Following bleaching, embryos were re-incubated in amplification buffer and incubated with a new subset of fluorescent hairpin amplifiers corresponding to additional target genes. This cycle of amplification, imaging, bleaching, and re-staining was repeated iteratively, enabling visualization of up to ten genes within the same embryo. Images generated from different sequential imaging rounds were subsequently aligned computationally using the image registration procedure described below.

### Chemical Bleaching and Photobleaching Procedure for Sequential Imaging

Following each imaging round, fluorescent signal was removed using a combination of chemical bleaching and limited photobleaching to prevent fluorescence carryover between sequential imaging cycles. All solution exchanges during bleaching and washing steps were performed directly within the Bioptechs FCS2 imaging chamber by pipetting solutions through the chamber inflow inlet while excess liquid exited through the outlet port.

For preparation of the MCPBA bleaching stock solution (0.1 M), 0.045 g meta-chloroperoxybenzoic acid (MCPBA) was dissolved in 2 mL of 97% ethanol. The stock solution was protected from light and stored at 4°C for up to 36 h before use. Immediately before bleaching, a fresh working solution was prepared by mixing 12 µL of 0.1 M MCPBA stock solution with 988 µL PBS. Embryos were then incubated in the MCPBA working solution within the imaging chamber for 1 h at room temperature.

Following MCPBA treatment, embryos were washed once with PBT through chamber-mediated fluid exchange and subsequently exposed to reducing solution. The reducing solution (1 M sodium thiosulfate) was prepared by dissolving 1.24 g sodium thiosulfate in 5 mL PBS and stored at 4°C. Embryos were incubated in reducing solution for at least 10 min to neutralize residual oxidizing agent and facilitate removal of remaining fluorophore signal.

After reducer incubation, embryos were washed extensively with PBT followed by PBS through repeated fluid exchange within the chamber before subsequent amplification and staining cycles. In most cases, chemical bleaching efficiently removed fluorescent signal, and additional photobleaching was either unnecessary or limited to approximately 1–2 min for embryos exhibiting persistent residual fluorescence. Photobleaching was performed directly on the confocal microscope using high-intensity laser exposure.

This combined chemical bleaching and limited photobleaching strategy enabled repeated cycles of staining, imaging, and bleaching while preserving embryo morphology, positional stability, and probe integrity throughout sequential imaging experiments.

### Probe Design and Sequences

To detect spatial and temporal expression patterns during sequential imaging experiments, gene-specific HCR probes were designed against both exonic and intronic regions of selected Tribolium segmentation genes. Exonic probes were used to visualize mature mRNA localization, while intronic probes were used to detect nascent transcriptional activity within nuclei.

The probe sets targeted the following genes: *eve, run, odd, hb, Kr, mlpt, gt, svb, cad,* and *en*. Intronic probe sets were generated for selected genes including *Kr, mlpt,* and *svb* to distinguish active transcription sites from accumulated cytoplasmic transcripts.

All probe sequences were synthesized for hybridization chain reaction (HCR)-based fluorescent detection and were compatible with the sequential imaging workflow described above. Complete probe sequences used in this study are provided in Supplementary Materials and Methods.

### Image Alignment of Sequential Fluorescence Images

Imaging the same embryo across multiple sessions introduces spatial misalignments that must be corrected before any quantitative comparison can be made. Differences in embryo orientation on the slide, positional drift between mounting sessions, minor variations in effective magnification, and small amounts of tissue deformation between fixation rounds all contribute to misregistration between successive images. A pixel at a given coordinate in one session does not necessarily correspond to the same anatomical location in another. All images of a given embryo must therefore be brought into a common spatial coordinate frame prior to analysis.

To this end, we developed a fully automated registration pipeline. The DAPI channel from the first imaging round was processed and used as a fixed structural reference. For each subsequent round, the DAPI image was normalized to correct for intensity variation, smoothed to reduce high-frequency noise, and thresholded to isolate the embryo signal. Morphological cleaning operations were then applied to remove small spurious foreground regions and fill internal holes, yielding a soft embryo mask for that round. Each round’s mask was registered to the mask of the first round by estimating a similarity transform that corrected for in-plane translation, modest differences in effective magnification, and minor rotational offsets. Once the transform was estimated from the DAPI channel of each round, it was applied identically to all corresponding gene expression channels, ensuring a consistent spatial coordinate system across genes imaged in different rounds. The DAPI channel itself was excluded from the final registered output.

Alignment quality was evaluated quantitatively by computing the DAPI mask overlap score between the reference and each subsequent round, which increased consistently after registration across all embryos. DAPI overlay images were then inspected visually to confirm that embryo boundaries and internal landmarks were in agreement after registration. Once alignment was confirmed, the registered gene expression channels were saved for downstream analysis.

## Spatial Gene Expression Extraction from Sequential Images

### 1. Region of interest and mesoderm exclusion

Once all imaging sessions for a given embryo are aligned to a common coordinate frame, gene expression signals are extracted from the fluorescence images. A region of interest (ROI) spanning the full dorso-ventral width of the embryo is drawn manually in ImageJ. The ventral midline region, which corresponds to the mesoderm, is explicitly excluded from the ROI. The mesoderm can exhibit non-specific background fluorescence that does not reflect ectodermal gap gene activity, so including it would confound the extracted intensity profiles. Only the ectodermal signal along the embryo body is retained for quantification.

### 2. Anterior-posterior intensity profiles

Within the defined ROI, mean fluorescence intensity is computed in narrow bins along the anterior-posterior (AP) axis. This yields a one-dimensional spatial profile *I*_*g*_(*x*) for each gene *g*, where *x* denotes position along the AP axis. The axis is normalized to the interval [0,1], with *x* = 0 corresponding to the anterior end and *xx* = 1 to the most posterior end. This normalization places all embryos on a common spatial coordinate regardless of their absolute size or orientation, allowing direct comparison across imaging sessions and individuals.

Intensity values are then normalized along the y-axis to the [0,1] range on a per-gene, per-embryo basis, by dividing each profile by its maximum observed value. This removes differences in absolute fluorescence arising from variation in probe efficiency, laser power, or staining batch between sessions, and allows the expression patterns of multiple genes to be displayed on a common intensity axis within a single plot. Optional Savitzky-Golay smoothing is applied to reduce high-frequency noise from local intensity fluctuations without distorting the positions or relative heights of expression domains.

### 3. Expression blocks representation

A compact block representation of spatial expression is constructed for each gene to complement the line-plot profiles. Each gene is assigned to a horizontal row. Within each row, the normalized expression intensity along the AP axis is rendered as a continuous color gradient, ranging from white at zero expression to a gene-specific saturated color at maximum expression. Because raw fluorescence intensity values, even after normalization, can appear perceptually flat when rendered as a linear color gradient, a perceptual enhancement algorithm is applied to improve the legibility of sharp expression boundaries and stripe structure. The algorithm boosts colors at positions where expression rises sharply and leaves low-level background regions faint.

For each AP position *x*, a local contrast measure *r*(*x*) is computed as the absolute difference between the normalized intensity 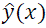 and the minimum intensity within a symmetric neighborhood window of half-width *w* = 0.05 (*in normalized AP units*):

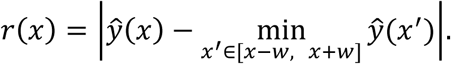

A perceptual contrast weight *P*(*x*) is then computed by thresholding *r*(*x*) against a saturation level *r_sat_* = 0.175:

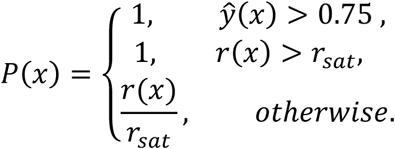

The final perceptual intensity *I*_perc_(*x*) is a convex combination of the raw normalized intensity and the contrast weight:

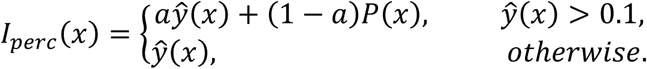

where *a* = 0.3 is a fixed blending coefficient. The condition 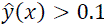 ensures that background pixels close to zero are not enhanced and remain white. This non-linear mapping amplifies local contrast at expression boundaries, making stripe positions and expression domain edges clearly visible in the block representation

## Temporal gene expression profiling

Temporal expression profiling was designed to capture how gap gene activity evolves across developmental stages without requiring re-imaging of the same embryo over time. Instead, cohorts of embryos were fixed, stained, and imaged at defined developmental stages, yielding a cross-sectional view of how mean expression changes across the full 24-stage series. All measurements are gene-specific: each gene was quantified independently from a set of embryos, and no intensity value is shared between genes. For each gene and each stage, five biologically independent embryos were used, giving a total of 24 × 5 = 120 embryos per gene.

Gap gene expressions are restricted to the posterior end of the embryo especially at early stages, so the anterior region contains little to no genuine gene signal. The anterior region therefore serves as a local background reference, capturing non-specific fluorescence, probe background, and imaging noise present across the embryo. For each embryo, two ROIs were drawn manually in ImageJ: one covering the posterior region and one covering the anterior region. In both cases, the mesoderm was excluded. The mean pixel intensity within each ROI was then extracted for the fluorescence channel corresponding to the gene of interest using the ImageJ Measure function. The mean anterior intensity 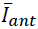 is subtracted from the mean posterior intensity 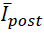 to yield a background-corrected signal:

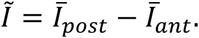

The corrected intensity values were normalized on a per-gene basis to put all genes on a common amplitude axis and the five normalized values at each developmental stage were then averaged to yield a single representative mean intensity per stage. The resulting temporal profile was visualized by plotting the stage label (ordered from 1·1 to 8·3) on the horizontal axis against the normalized mean intensity on the vertical axis.

## Simulation of Intronic and Exonic Expression Along the Anterior-Posterior Axis

To examine how a spatially travelling gene expression wave manifests as a temporal pulse when observed at a fixed posterior location, we simulated a Gaussian expression pulse moving at constant speed from posterior to anterior along the normalized AP axis *Z* ∈ [0,1], with *Z* = 0 corresponding to the anterior end and *Z* = 1 to the posterior end. The domain was discretized into spatial points, and the simulation was run for a total time *T* = 5.

The intronic signal *x*(*Z*, *t*) was prescribed as a Gaussian pulse of amplitude *A* and width σ travelling at a constant speed *v*:

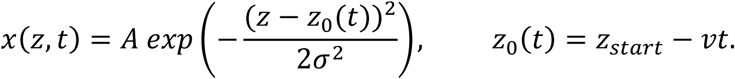

The pulse center starts at *Z_start_* = 1 + 3σ, placing it just outside the posterior boundary at *t* = 0, so that it enters the domain smoothly and sweeps towards the anterior over time. The exonic signal *y*(*Z*, *t*) was coupled to the intronic signal through a first-order production-decay equation:

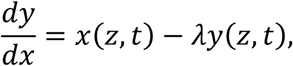

where λ is the decay rate. This equation was integrated forward in time using the Euler method. Parameter values used were *A* = 1, σ = 0.08, *v* = 0.2, and λ = 1.

The time courses of both the intronic and exonic signals were recorded at a fixed observation point at the posterior end of the domain, **Z*_*o*b*s*_* = 1. This mimics the temporal expression profiling experiment, where mean fluorescence intensity is measured at the posterior end of each embryo across developmental stages. As the simulated wave sweeps anteriorly past **Z*_obs_*, both signals produce a transient temporal pulse, directly demonstrating that the temporal expression profile recorded at the posterior is a consequence of a travelling spatial wave passing through that fixed location. To allow direct visual comparison of the intronic and exonic time courses, the exonic signal was rescaled at each time point so that its maximum matched that of the intronic signal, preserving the shape and timing of the profile while removing the amplitude difference introduced by the integration.

GitHub: sweeping_pulse_spacetime.m

## Single-Gene Speed Regulation Simulation Under Three Gradient Regimes

To illustrate how the speed regulation model generates spatiotemporal gene expression patterns, we simulated a single-gene system in which a morphogen gradient controls the local rate at which each cell accumulates transcriptional phase. The AP axis was represented as a normalized spatial domain, and the simulation was run from *t* = 0 to *t* = 5.

Each cell *j* accumulates phase *y*_*j*_ (*t*) at a rate set by the local morphogen concentration

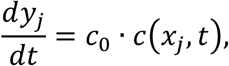

where *c*_0_ = 0.3 is a baseline speed multiplier. The gene switches on when the accumulated phase enters the expression window defined by the ON and OFF thresholds *y_on_*= 0.3 and y*_off_* = 0.5. Rather than a hard binary switch, the gene state was modelled as a smooth Gaussian function of phase, centered at the midpoint of the ON/OFF window with width 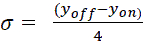. This yields a continuous intronic signal *x_j_* (*t*) that rises and falls smoothly as the cell’s phase passes through the expression window. The exonic signal *e_j_* (*t*) was coupled to the intronic signal through a first-order decay equation 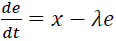, with decay rate λ = 1, and integrated forward using the Euler method.

Three forms of the morphogen gradient *c*(*x*, *t*) were used to represent the three modes of the speed regulation model: the gradient-based, the wavefront-based and the graded wavefront. In the gradient-based mode, the morphogen is a static linear ramp that increases from *c_min_* = 0 at the anterior to *c_max_* = 1 at the posterior and does not change over time. Posterior cells therefore accumulate phase faster and enter the expression window earlier, producing sequential waves of gene expression that sweep from posterior to anterior. This mode corresponds to long-germ patterning, where AP fates are specified primarily in the blastoderm under a smooth, non-retracting gradient.

In the wavefront-based mode, the morphogen is a sharp step function whose boundary starts at the anterior and retracts posteriorly at speed *v* = 1.5 × 0.12. Cells to the right of the boundary are exposed to *c_max_* and accumulate phase rapidly; cells to the left receive *c_min_* and do not progress. As the boundary sweeps posteriorly, it sequentially releases cells from high morphogen exposure, freezing their expression state into stable spatial domains. This mode corresponds to short-germ patterning, where a retracting wavefront stabilizes AP fates in the germband. In the graded wavefront mode, the step is replaced by a smooth linear transition zone of finite width Δ*x_step_* that also retracts posteriorly at the same speed. This intermediate regime combines features of both modes. It generates sequential expression waves like the gradient-based mode, but the retracting boundary progressively freezes the pattern like the wavefront-based mode. It corresponds to intermediate-germ patterning, where both blastoderm and germband contribute to AP fate specification.

GitHub: speed_regulation_three_gradients.m

## Effect of Gradient Shape on Expression Domain Widths (Fig. 7A)

The shape of the morphogen gradient affects the spatial distribution of gene expression domains along the AP axis. To show this, we simulated a single-gene speed regulation system under two gradient forms of the gradient-based mode. Phase accumulation at each cell position followed:

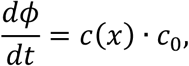

with baseline speed multiplier *c*_0_= 0.3. The gene state at each position was computed as a Gaussian function of the accumulated phase *d* and only the intronic signal was tracked in this simulation.

The first gradient was a linear speed gradient 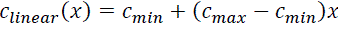. The second was a concave (power-law) speed gradient 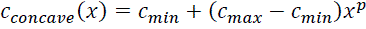, with *p* = 0.5. Both used *c_min_* = 0 and *c_max_* = 1 and were static in time. The two simulations were run in parallel to allow direct comparison of domain widths across the AP axis.

GitHub: gradient_shape_domain_widths.m

## Quantification of Space-to-Time Correspondence Using Levenshtein Fidelity

To quantify how faithfully the spatial ordering of gene expression domains along the AP axis corresponds to the temporal ordering of gene activation, we applied a sequence-based fidelity analysis to two gene groups separately: pair-rule genes (*eve*, *runt*, *odd*) and gap genes (*kr*, *mlpt*, *gt*). The analysis was performed across four developmental stages (3.3, 4.1, 4.3, and 5.2), with three spatial embryos per stage. For each embryo, the normalized spatial expression profile was windowed to a region of interest spanning the AP axis positions over which *eve* expression is visible. The temporal expression profile for the same gene was similarly windowed to the corresponding range of developmental stages. Within each windowed profile, a set of landmark points was selected along the x-axis for each gene and assigned a single-letter gene prefix. Each landmark corresponded to a local feature of the expression profile, such as a peak, a trough or a stripe boundary. The landmarks across all gene groups within an embryo were then pooled and sorted by their normalized x-coordinate, producing an ordered sequence of labelled landmarks.

The similarity between a spatial sequence and the temporal sequence was quantified using the Levenshtein edit distance. Only the gene-letter prefix of each landmark label was used in the comparison, so that the distance reflected the ordering of genes rather than the ordering of individual landmarks within a gene. The Levenshtein fidelity score *F* was defined as

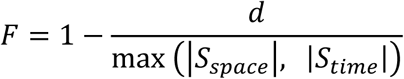

where *d* is the Levenshtein distance between the spatial sequence *S_space_* and the temporal sequence *S_time_*, and the denominator is the length of the longer of the two sequences. A fidelity of 1 indicates a perfect match between the spatial and temporal gene orderings, and a fidelity of 0 indicates no correspondence.

